# OLD sentinel: an abortive tRNase surveys phage replication and DNA defects in RecBCD-compromised cells

**DOI:** 10.64898/2026.07.10.737580

**Authors:** Anton Derzhaev, Jing Zhang, Alexey Gavrilov, Svetlana Belukhina, Aleksandr Shenfeld, Florence Depardieu, Baptiste Saudemont, Ilya Shamovsky, Vitaly Epshtein, Alina Demkina, Huaile Song, Olga Burenina, Mikhail Skutel, Maria Tikhomirova, Vadim Molodtsov, Konstantin Severinov, Evgeny Nudler, David Bikard, Chengyuan Wang, Artem Isaev

## Abstract

OLD, an abortive immunity protein from prophage P2, consists of an ABC ATPase sensor and a TOPRIM nuclease effector — a core architecture shared by a large protein family, including components of antiphage systems Gabija, PARIS, Septu, and Lamassu. OLD was originally identified for its lethality in *recB*-deficient cells and inhibition of bacteriophage λ infection, but the mechanisms governing its activation have remained elusive. Here, we present the cryo-EM structure of an inactive OLD tetramer and show that destabilization into dimeric form opens the TOPRIM catalytic site, stimulating tRNA cleavage. This activity arrests translation, a phenotype rescued by phage-encoded tRNAs. We demonstrate that OLD activation is not strictly RecBCD-dependent: OLD binds aberrant DNA structures in *recB*-deficient cells, but activation during infection requires recognition of single-stranded DNA hairpins at the phage replication origin. Collectively, our findings reveal how host and phage DNA processing factors create a complex landscape controlling OLD-mediated immunity.

## Introduction

The ever-ongoing evolutionary arms race between bacteria and their viral predators, bacteriophages, has driven the diversification of sophisticated defense and counter-defense mechanisms^1,2^. These bacterial defenses can be conceptualized as operating in distinct “layers”. The first layer comprises direct immunity systems — such as Restriction-Modification (RM) and CRISPR-Cas — which recognize invading phage DNA and eliminate it to protect the infected cell. However, this direct immunity often fails due to phage-encoded countermeasures like DNA modifications and anti-defense proteins (e.g., Acrs, Ocr)^3,4^. Consequently, a second layer of defense, known as abortive infection (Abi) systems, can act at later infection stages and often provide population-level protection by driving the infected cell into dormancy or death^5,6^. This outcome can be directly induced by the Abi effector or can result from the toxicity of phage proteins produced within the cell^6^.

Abortive infection (Abi) systems that actively induce cell-toxicity typically possess a modular architecture, comprising a sensor domain that detects phage infection and a toxic effector domain that targets essential host macromolecules^7^. Phage infection is recognized either directly, through the binding of specific phage proteins (triggers) to the sensor domain^8–10^, or indirectly, via phage-induced metabolic perturbations (inhibition of host transcription^11^, accumulation of dNMPs and free DNA ends upon host DNA degradation^12–14^, or the exhaustion of cellular nucleotide pools^15,16^). Immunity effector activation often relies on oligomerization or a conformational switch, a step that could be regulated by ATPase domains, integrating both types of signals.

Some immunity systems, such as PARIS and Panoptes, can specifically recognize viral anti-immune proteins (DNA mimics or signaling molecule sponges)^17–21^ as triggers of the abortive response. Multiple phages also encode inhibitors of the host nuclease-helicase RecBCD, a key enzyme in homologous recombination and dsDNA break repair^22–24^. RecBCD efficiently degrades linear dsDNA, and phages could expose dsDNA ends during their life cycle, however, the role of RecBCD in anti-phage defense remains controversial^25,26^.

It is assumed that phages encoding their own recombination systems (e.g., Exo, β of the phage λ), benefit from suppressing host recombination machinery using inhibitors like Gam^27^. Notably, immunity systems such as Ec48, Se72, AbpAB, DRT3 or the prophage P2 OLD, are activated in response to phage RecBCD inhibitors or in *recB*-deficient cells^28–32^. Hence, the RecBCD complex represents another focal point of phage–host conflict: it directly influences phage recombination and replication^33^, serves as a target for inhibition^34–36^, and controls activation of immunity systems. Yet the mechanism by which immunity systems surveil RecBCD activity remains undetermined.

OLD (for <u>o</u>vercoming <u>l</u>ysogenization <u>d</u>efect) was discovered by G. Sironi in 1968 as a phage P2 gene incompatible with lysogenization of recombination-deficient *Escherichia coli* host^30^. In subsequent work, it was shown that OLD kills *recBC* compromised cells and protects P2 lysogens against phage λ infection^37^. The isolation of escapers revealed that λ actively interferes with host recombination, a discovery that led to the description of Gam as an inhibitor of RecBCD^38^ and likely OLD trigger. In parallel, F. Brégégère demonstrated that OLD can be activated upon prophage λ induction. This activation requires both phage (O, P) and host (DnaB) proteins involved in the initiation of λ replication and results in damage to the host tRNAs^39–41^. In contrast, later works focused on testing OLD’s activity against DNA *in vitro*, leading to its characterization as an exonuclease^42–45^. Therefore, conflicting reports exist regarding the mechanisms of OLD activation and its target, an ambiguity that we aimed to resolve in this study.

Proteins from the OLD family are composed of ABC ATPase and TOPRIM nuclease domains—a modular architecture observed in a variety of phage immunity systems, in which the ABC domain acts as a sensor and/or conformational switch that controls the TOPRIM effector, which can act as a DNase or RNase. ATPases from the ABC superfamily are essential for energy-coupled processes including membrane transport, DNA repair, chromosome organization, and translation regulation^46^. A dimer of ABC domains is required to bind ATP at two composite sites, where ATP binding itself rather than hydrolysis represents the power stroke that drives the conformational change of the complex^47^. Within the ABC superfamily, OLD ATPases are often found in highly mobile genetic loci across prokaryotes, archaea, and some eukaryotes^46^.

Classification of OLD-like systems is rather complex^46,48^. While OLD is a standalone immunity system, a two-protein module composed of OLD (GajA) and UvrD-like helicase (GajB) has been called a Gabija system^49–53^; OLD can be associated with a retron reverse transcriptase that synthesizes msDNA–RNA hybrids, acting as phage SSB protein sensor^54–57^. ABC and TOPRIM domains can represent separate proteins, like in PARIS (that senses phage DNA mimic proteins), AbiL, and MADS systems^18,58,59^. Beyond this, ABC+TOPRIM module appear in multiple toxin–antitoxins and poorly described Abi systems^60–63^. In addition to TOPRIM, ABC ATPases can be coupled to diverse effector domains: in Septu and PD-T4-4 systems it is an HNH nuclease^64,65^, in PrrC and RloC it is HEPN tRNAse^66,67^, while in Menshen systems ABC domain functions as a conformational switch/activator coupled to diverse sensor/effector domains^68,69^. A distinct type of ABC domain with an extended coiled coil insertion is found primarily in DNA-repair complexes such as Rad50–Mre11 and SbcCD, and in Lamassu immunity system^70–72^. Furthermore, proteins with even longer coiled coil region and a hinge domain are involved in DNA compactization (Structural Maintenance of the Chromosome or SMC proteins) and gave rise to a Wadjet anti-plasmid immunity system^46,73^. Notably, OLD homologs can regulate DNA replication, as recently shown in archaea^74^. Collectively, this demonstrates that the OLD-like ABC+TOPRIM architecture can be adapted to a broad range of functions, oligomeric states, partner proteins, and input signals (DNA or protein), making it one of the most widespread frameworks in bacterial immunity.

Here, we describe the mechanisms of activation and defense for a prototypical OLD protein from prophage P2. A cryo-EM structure of OLD reveals a tetrameric architecture stabilized by specific inserts in the ABC domains, while the TOPRIM domains dimerize and are held in an inactive, closed conformation. Destabilization of the tetramer activates OLD, leading to opening of the TOPRIM catalytic cleft. We demonstrate that OLD cleaves host tRNAs both *in vivo* and *in vitro*, and that its anti-phage activity can be suppressed by supplying cleavage-resistant phage tRNAs. Finally, we show that OLD activation does not strictly rely on RecBCD activity; instead, OLD is triggered by aberrant DNA structures that accumulate in the host terminus upon RecBCD deficiency, or by ssDNA hairpins in the phage replication origin. In addition to RecBCD, this activity is modulated by both phage recombination system and the host ssDNA hairpin-cleaving enzyme SbcCD.

## Results

### Recombination-related loci of various phages encode OLD triggers and inhibitors

OLD protein is encoded in an immunity hotspot region of the P2 prophage genome (Figure 1A), and provides defense against phage λ infection^75^. We cloned P2 OLD under its native promoter and confirmed that it confers robust protection against λ, as well as members of the *Hendrixvirinae* and *Dhillonvirus* groups, not previously reported to be OLD-sensitive (Figure 1B). A weak defense phenotype was also observed against phage T7, which encodes a RecBCD inhibitor gp5.9, and against the T5-like phage Bas34^76^. OLD activity required intact ABC ATPase and TOPRIM domains, since mutations in the ATPase H-loop (H332A) or TOPRIM catalytic site (E402A) abolished the defense^44^ (Figure S1A). We have also constructed an anhydrotetracycline (aTc)-inducible pFR_OLD plasmid that provided mild protection without induction (termed OLD_Low_ in later experiments) and complete defense upon induction (Figure S1A).

**Figure 1.**
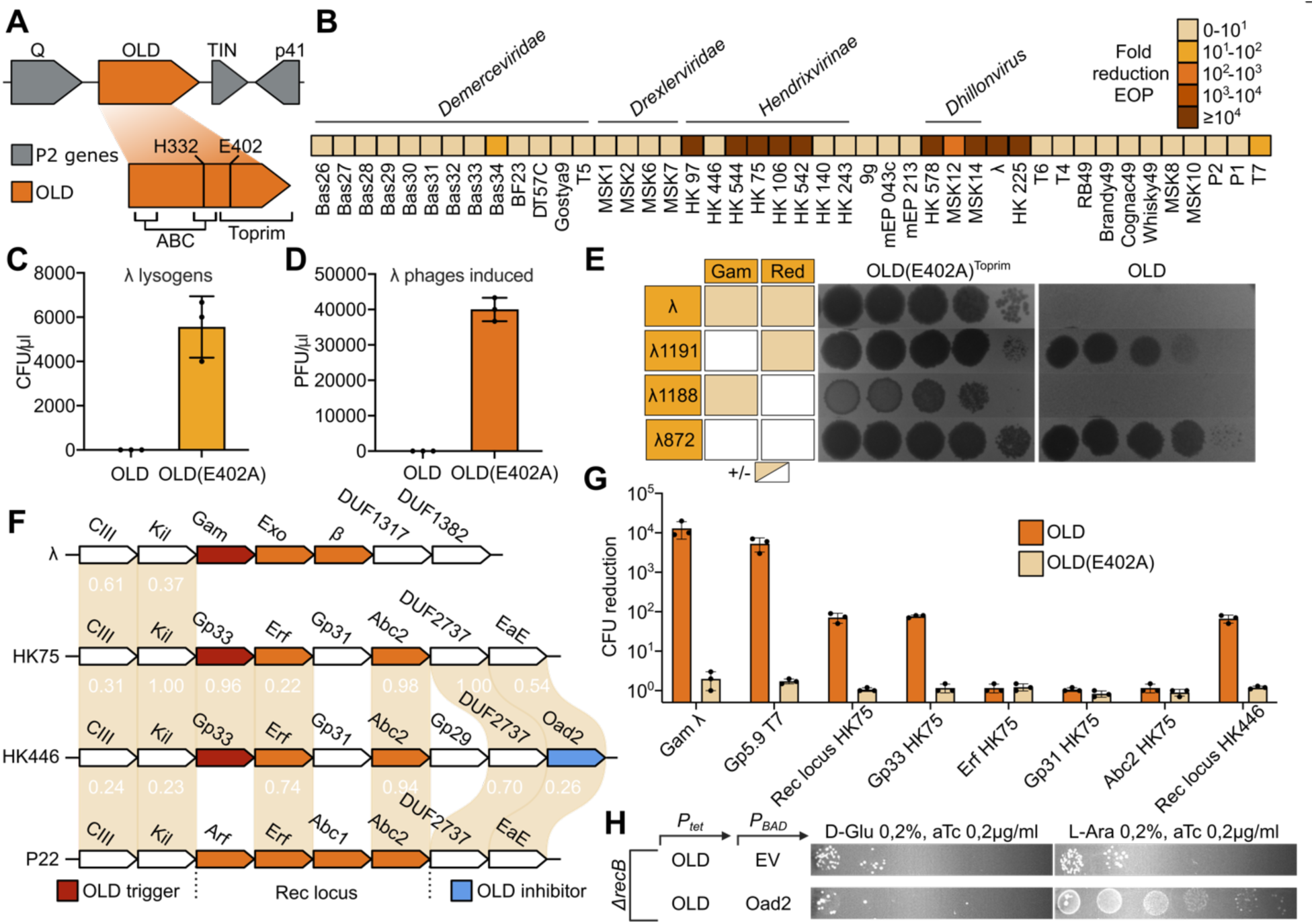
Characterization of OLD defense system. Discovery of a novel trigger and inhibitor. **(A)** Schematic of the P2 prophage defense locus encoding the OLD and Tin systems. OLD is composed of an N-terminal ABC ATPase domain and a C-terminal TOPRIM nuclease domain. **(B)** Defense spectrum of OLD against diverse *E. coli* phages, measured by efficiency of plating (EOP). **(C)** Efficiency of lysogenization of phage λ_ts_(CmR) in cells expressing OLD or the TOPRIM catalytically inactive mutant OLD(E402A). **(D)** Efficiency of prophage λ_ts_(CmR) induction from сells expressing OLD or OLD(E402A). **(E)** EOP of phage λ and its derivatives λ1191(*gam^-^),* λ1188 (*beta^-^*) and λ872 (*gam^-^ exo^-^ beta^-^*), demonstrating that loss of Gam enables escape from OLD-mediated defense. **(F)** Genetic organization of CIII-regulated recombination-related loci in phages λ, HK75, HK446, and P22. **(G)** Toxicity assay results showing colony-forming unit (CFU) reduction upon expression of phage proteins in OLD or OLD(E402A) cells. Rec locus includes gp33, Erf, gp31 and Abc2 encoding genes. **(H)** Toxicity of inducible OLD in Δ*recB* cells and rescue by co-expression of Oad2.

OLD activity was not restricted to the phage lytic cycle, as measured by efficiency of plating (EOP), but also inhibited λ lysogenization and prophage induction (Figures 1C and 1D). This indicates that OLD could sense λ-specific triggers produced both during early infection (lysis/lysogeny decision stage) and during active replication cycle, which is in contrast to the GajAB Gabija defense system that has been reported to be inactive during λ-like prophage D3 induction^77^.

To validate prior reports that λ mutants lacking the RecBCD inhibitor Gam evade OLD defense^37^, we employed three mutant phages: λ1191 lacking Gam (*gam^-^ red^+^)*, λ872 lacking Gam and a recombination system (*gam^-^ red^-^*) and λ1188 lacking only recombination proteins (*gam^+^ red^-^*) as a control^78^. EOP assays demonstrated that λ1191 and λ872, but not λ1188, formed plaques on OLD^+^ cells (Figure 1E), albeit forming smaller-sized plaques, suggesting that Gam is required to trigger OLD. Consistent with this, heterologous expression of RecBCD inhibitors λ Gam or T7 gp5.9 was toxic to OLD^+^ cells in the absence of phage infection (Figure 1G), confirming that RecBCD inhibition by viral proteins is sufficient to activate OLD. In accordance with these results and with previous observations^30^, OLD expression in *recB*-deficient host was toxic (Figure S1B).

Some OLD-sensitive phages, such as HK75, lack known RecBCD inhibitors. To predict possible OLD triggers associated with these phages, we aligned their genomes with λ and P22 (Figure 1F), encoding distinct types of homologous recombination loci containing known RecBCD manipulation genes. In all phages, these loci are flanked by the CIII-like repressor gene. In phage λ, an exonuclease Exo and a single-strand annealing protein (SSAP) β are coupled with RecBCD inhibitor Gam; phage P22 encodes SSAP Erf and an array of poorly-studied proteins assisting in recombination and altering RecBCD activity (Arf, Abc1, Abc2)^79^. In HK75 and HK446 the corresponding operon encodes a mixture of P22-like (Erf and Abc2) and uncharacterized (Gp31, Gp33) proteins. To identify genes responsible for OLD activation, we cloned the full HK75 recombination locus (Gp33, Erf, Gp31, Abc2), as well as its individual genes, and found that Gp33 expression is sufficient to trigger OLD toxicity to the same extent as the full recombination locus. Considering that other OLD triggers are RecBCD inhibitors we hypothesize that Gp33 could represent a novel RecBCD inhibitor or manipulator.

Intriguingly, phage HK446 evaded OLD-mediated defense (Figure 1B) despite encoding the Gp33. Moreover, expression of the HK446 recombination locus also triggered OLD toxicity (Figure 1G). Thus, we hypothesized that in addition to OLD trigger, HK446 encodes an OLD inhibitor. To identify it, we constructed a randomized plasmid library from the HK446 genome and screened for constructs that could rescue OLD toxicity in *ΔrecB* cells (Figure S1C). This screen identified multiple plasmids carrying inserts with gene *gp27*, which we confirmed to be sufficient to inhibit OLD toxicity in *recB*-deficient host and anti-phage defense (Figures 1H and S1D). Following a previously proposed nomenclature^80^, we designate this protein Oad2 (OLD anti-defense 2). Notably, *oad2* (*gp27*) gene is encoded close to the OLD trigger *gp33*, suggesting a hotspot of anti-defense genes associated with phage homologous recombination loci that activate various immunity systems^29,81^. OLD-sensitive phage HK75 also encodes a homolog of *oad2* with 54% amino acid identity. The fact that HK75’s homolog is not sufficient to inhibit OLD defense suggests a highly specific interaction required to achieve immunity suppression, a signature of the ongoing arms race at the evolving protein/inhibitor interface.

### OLD activation results in host tRNA cleavage

To elucidate the mechanism of OLD-mediated defense, we started by confirming its abortive phenotype: at high multiplicity of infection (MOI) phage λ does not lyse OLD^+^ culture, yet it causes early cell growth arrest not observed in the control culture (Figures 2A and S2A). Since conflicting reports have characterized OLD as either a DNase or a tRNAse^80,82,83^, we sought to determine its primary target *in vivo*. As shown previously, tRNA cleavage and translation arrest is associated with a specific DNA-compaction phenotype^84^. Microscopic analysis revealed that cells co-expressing OLD and its trigger protein, Gam, exhibited a compacted nucleoid morphology (Figure 2B) with no evidence of DNA degradation in either microscopy, total DNA purification or TUNEL assay (Figures 2B, S2B and S2C). This suggests that a primary *in vivo* OLD target is associated with translational machinery. At the same time, total RNA purified from *ΔrecB* cells after OLD expression demonstrated a cleavage pattern reminiscent of tRNA fragmentation (Figure 2C). Consequently, we performed tRNA sequencing (tRNA-seq) of cells expressing OLD or a GFP control following Gam induction. Quantification of intact tRNA reads revealed a significant depletion (>2-fold) of specific tRNAs (Figure 2D), including Thr(UGU), Gly(UCC), fMet(CAU), Val(UAC), and Pro(UGG). Mapping of 5’ and 3’ termini pinpointed precise endonucleolytic cleavage events, predominantly after nucleotides 27, 28, or 29, with the exact site varying by tRNA isoacceptor (Figure S2D). A subset of tRNAs also exhibited a secondary cleavage pattern consistent with sequential processing, and anticodon loop amputation, as reported in a parallel study^85^. To confirm RNA-seq observations, we performed a Northern blot analysis with the specific probes complementary to Thr(UGU) tRNA that demonstrated complete tRNA depletion upon OLD expression in *ΔrecB* cells (Figure S2E).

**Figure 2.**
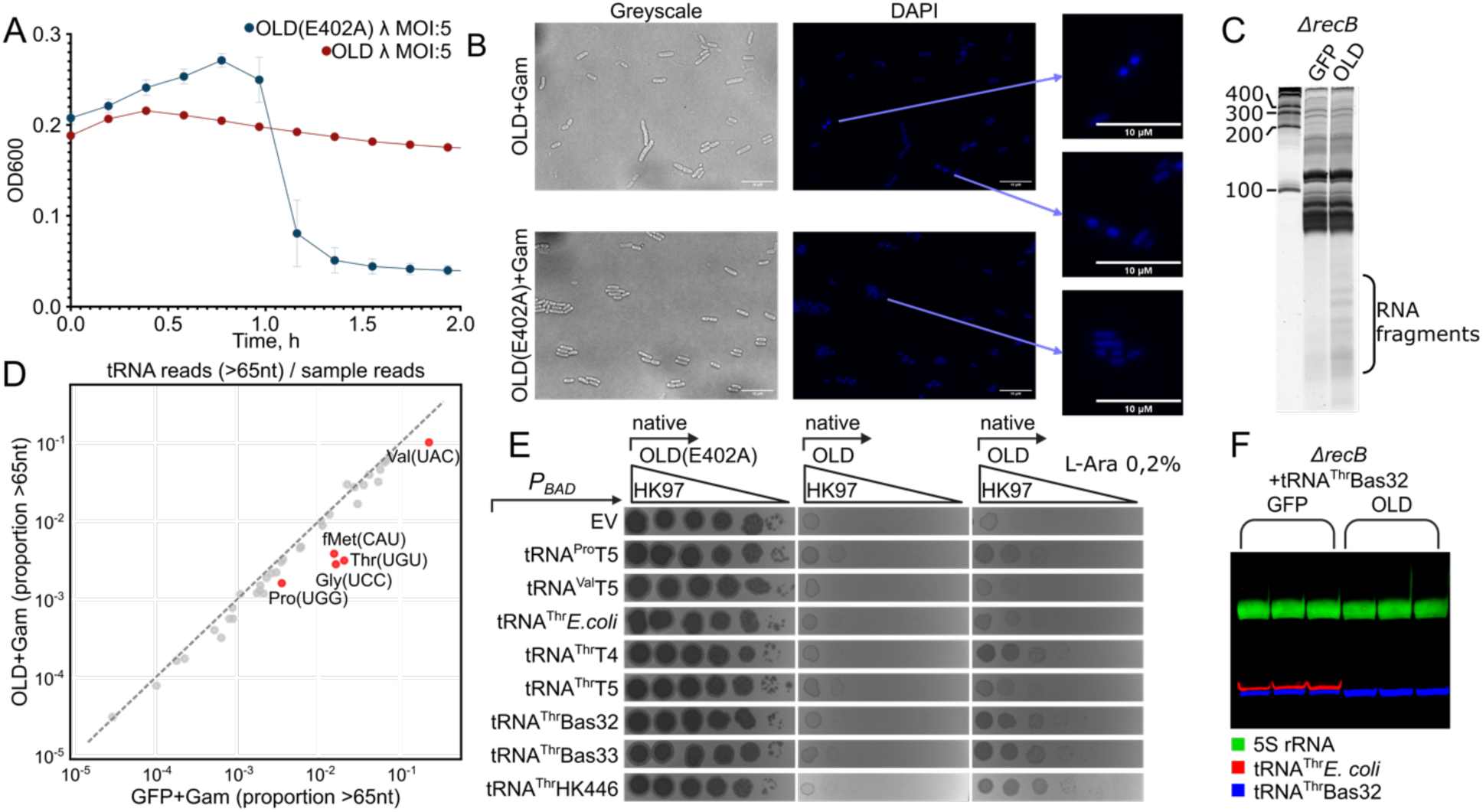
OLD is an abortive infection system that targets host tRNAs. **(A)** Growth curves showing that at high multiplicity of infection (MOI), OLD-expressing cultures cease growth before phage lysis, consistent with an abortive infection phenotype. **(B)** DAPI staining of cells co-expressing OLD and Gam reveals compacted nucleoids (blue), indicative of translation arrest. Enlarged insets highlight distinct nucleoid morphologies in OLD + Gam (top and middle) compared to the catalytically inactive OLD (E402A) + Gam control (bottom). Images were taken 90 min post-induction. **(C)** Total RNA extracted from *ΔrecB* OLD culture demonstrates signs of fragmentation on denaturing (7M urea) PAGE. **(D)** Scatter plot showing the proportion of reads covering >65 nt of each tRNA relative to total reads in GFP/Gam (x-axis) versus OLD/Gam (y-axis) samples. Data from three biological replicates per condition were aggregated. Selected tRNAs showing the strongest depletion or enrichment (>2fold) are highlighted in red, while remaining tRNAs are shown in grey. The dashed gray line indicates X = Y. **(E)** Efficiency of plating (EOP) assay demonstrates that expression of viral tRNAs partially restores plaque formation of OLD-sensitive phage HK97. **(F)** Northern blot analysis of tRNA Thr(UGU) stability in Δ*recB* cells expressing OLD or GFP control. Endogenous *E. coli* tRNA Thr(UGU) is degraded, while phage Bas32 tRNA Thr(UGU) remains intact, confirming resistance to OLD cleavage. Hybridization was performed step by step with *E. coli* anti- tRNA Thr(UGU) probe (Cy3), followed by Bas32 anti- tRNA Thr(UGU) probe (Cy5) and finally with *E. coli* anti- 5S rRNA probe (FITC).

To functionally validate tRNA as both necessary and sufficient target associated with OLD toxicity, we performed a tRNA rescue experiment. It was recently demonstrated that viral tRNAs accumulate mutations making them insensitive to cleavage by abortive infection tRNAses^18,86^. Therefore, we have tested whether expression of Thr(UGU) tRNAs from *E. coli* and several phages (T4, T5, Bas32, Bas33, and HK446), along with Val(UAC) and Pro(UGG) tRNAs from phage T5, could inhibit OLD mediated anti-phage defense. While *E. coli* Thr(UGU) tRNA expression had no effect, viral Thr(UGU) tRNAs rescued infection of OLD-sensitive HK97, HK542 and Bas34 phages to a varied extent (Figures 2E, S3A, S3B and S3C). Expression of Thr(UGU) tRNAs from T4, Bas32, Bas33 and HK446 provided strong counter-defense, restoring plaque formation to near-wild-type levels. In contrast, Thr(UGU) tRNA from T5 conferred only partial rescue, suggesting that phage tRNAs possess varying degrees of intrinsic resistance to OLD cleavage.

To determine whether phage tRNAs function as competitive inhibitors (by binding to OLD and blocking its activity against host tRNAs), or whether they supplement cleaved host tRNAs without compromising OLD activity, we monitored the stability of host and viral tRNAs in one assay. Northern blot analysis of *ΔrecB* cells expressing OLD and supplemented with Bas32 Thr(UGU) tRNA revealed that the Bas32 tRNA remained stable, while endogenous *E. coli* Thr(UGU) tRNA was degraded to the same extent as in control lacking Bas32 tRNA (Figures 2F and S2E). These results demonstrate that phage-encoded tRNAs do not compromise OLD activity but rather supplement the tRNA pool, functionally replacing cleaved host tRNAs, confirming tRNA as OLD’s primary *in vivo* target. Whereas the expression of viral Thr(UGU) tRNA alone was insufficient to rescue OLD toxicity in Δ*recB* cells, neither to inhibit defense against phage λ (Figures S3D and S3E). This suggests that a full set of tRNA targets must be supplemented to completely inhibit OLD toxicity, as recently shown for the PARIS system^87^.

### Structure of the inactive OLD tetrameric complex

To elucidate the molecular mechanism by which OLD recognizes and cleaves tRNA, purified recombinant OLD protein was subjected to cryo-electron microscopy analysis, yielding a final reconstruction at 3.1 Å resolution (Figure S4 and Table S1). This map was of sufficient quality to build an atomic model.

The OLD protein consists of an N-terminal ABC ATPase domain and a C-terminal TOPRIM domain. Between these, an insertion within the N-terminal ABC ATPase region - termed the dimerization region - was identified (Figure 3A). Consistent with previously reported members of the OLD family nucleases, such as GajA^50^ and Eco8-OLD^54,55^, P2-OLD eluted as a tetramer in gel-filtration experiments and adopted a tetrameric architecture in the resolved structure (Figures 3A, S4B and 3B). The P2-OLD tetramer is assembled from two homodimers arranged in a tail-to-tail orientation, which are held together in a relatively tight embrace. However, unlike the GajA tetramer, where tetramerization relies primarily on interactions between ABC ATPase domains, the two homodimers in OLD associate via the dimerization region to form the complete tetramer (Figures 3A and S5).

**Figure 3.**
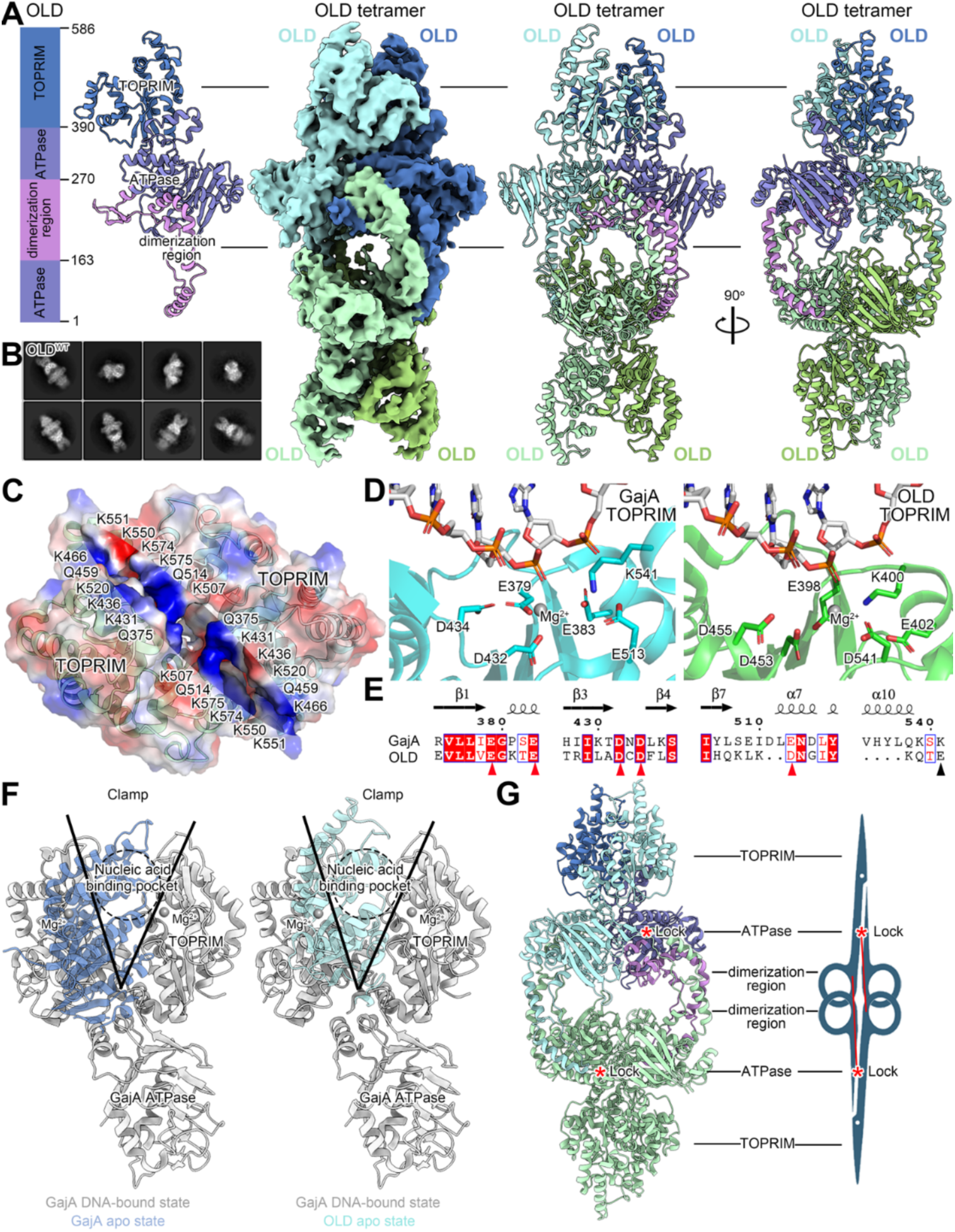
Cryo-EM structure of the autoinhibited OLD tetramer. **(A)** The overall structure of OLD tetramer. Schematic of the domain organization, the cryo-EM density map and ribbon diagrams of an OLD tetramer in two orthogonal views. The color scheme in the protomer is consistent in all panels. **(B)** 2D class average from cryo-EM analysis of OLD^WT^. **(C)** Enlarged top view of TOPRIM domain dimer within the OLD tetramer (surface colored by electrostatic potential). **(D)** Structure comparison of the predicted nuclease active sites in the TOPRIM domains of OLD and GajA. The OLD TOPRIM–DNA model (right), derived by fitting to the PDB 8WY5 template, is aligned with the experimental GajA–DNA structure (left). **(E)** Sequence alignment of the nuclease active site regions in OLD and GajA TOPRIM domains, key catalytic residues are indicated by red triangles. **(F)** Structural comparison of the OLD protomer with GajA in its DNA-bound state (PDB: 8WY5) and apo state (PDB: 8X5I) **(G)** Model of the OLD tetramer in the autoinhibited state. The binding site at the interface between dimerization and ATPase domains is marked with a red asterisk (*).

In the overall structure, the ATPase domains occupy a central position, forming a shoulder-like scaffold. The TOPRIM domains are located at the poles of the complex, linked by the ATPase domain (Figure 3A). Two neighboring TOPRIM domains create a channel for nucleic acid binding and cleavage, whose surface is enriched with basic residues (Figure 3C). At the center of the TOPRIM channel lies a conserved active site for nucleic acid cleavage. Although repeated attempts to obtain a substrate-bound complex were unsuccessful, the structural model reveals that conserved residues - including D455, D453, D541, E402, E398, and K400 - are positioned directly within the catalytic pocket (Figures 3D and 3E). Among these, the E402 residue has been confirmed by our data and previous reports to be critical for nuclease activity^44^. We performed a sequence-based structural alignment to compare the relative positions of the TOPRIM domains in the apo state of GajA, in the dsDNA-bound state of GajA, and in our tetrameric OLD structure (Figure 3F). The analysis revealed that the OLD tetramer resembles the apo state of GajA, with the channel formed between the two TOPRIM domains being too narrow to accommodate DNA or tRNA binding (13.5 Å distance between E398 residues of two protomers), suggesting an autoinhibited conformation (Figure 3G).

Interestingly, our findings suggest that the dimerization region in the OLD tetramer likely plays a critical role in both stabilizing the tetrameric structure and restricting the conformational switch of the TOPRIM dimer. From a structural perspective, this region not only reinforces tetramer stability through direct in-ter-subunit interactions but also engages with the ATPase domains, effectively acting as a molecular lock that secures the overall conformation of the tetramer (Figure 3G).

### OLD activation requires destabilization of the tetramer

The dimerization region of OLD adopts a comma-like shape composed primarily of several α-helices (Figure 4A). Within the tetramer, the dimerization region from each protomer in one OLD dimer interacts with the corresponding dimerization region from the opposite OLD dimer (Figure 3G), forming a head-to-tail binding arrangement between two pairs of dimerization regions (Figures 4A and 4B). A detailed zoom- in view revealed a set of hydrophobic residues (L170, I177, F185, I192, L196, L229, and F231) in the "body" of the comma-shaped region engaged in hydrophobic interactions with corresponding residues from the dimerization region of the opposing protomer, constituting the primary stabilizing interface of the tetramer (Figure 4B).

**Figure 4.**
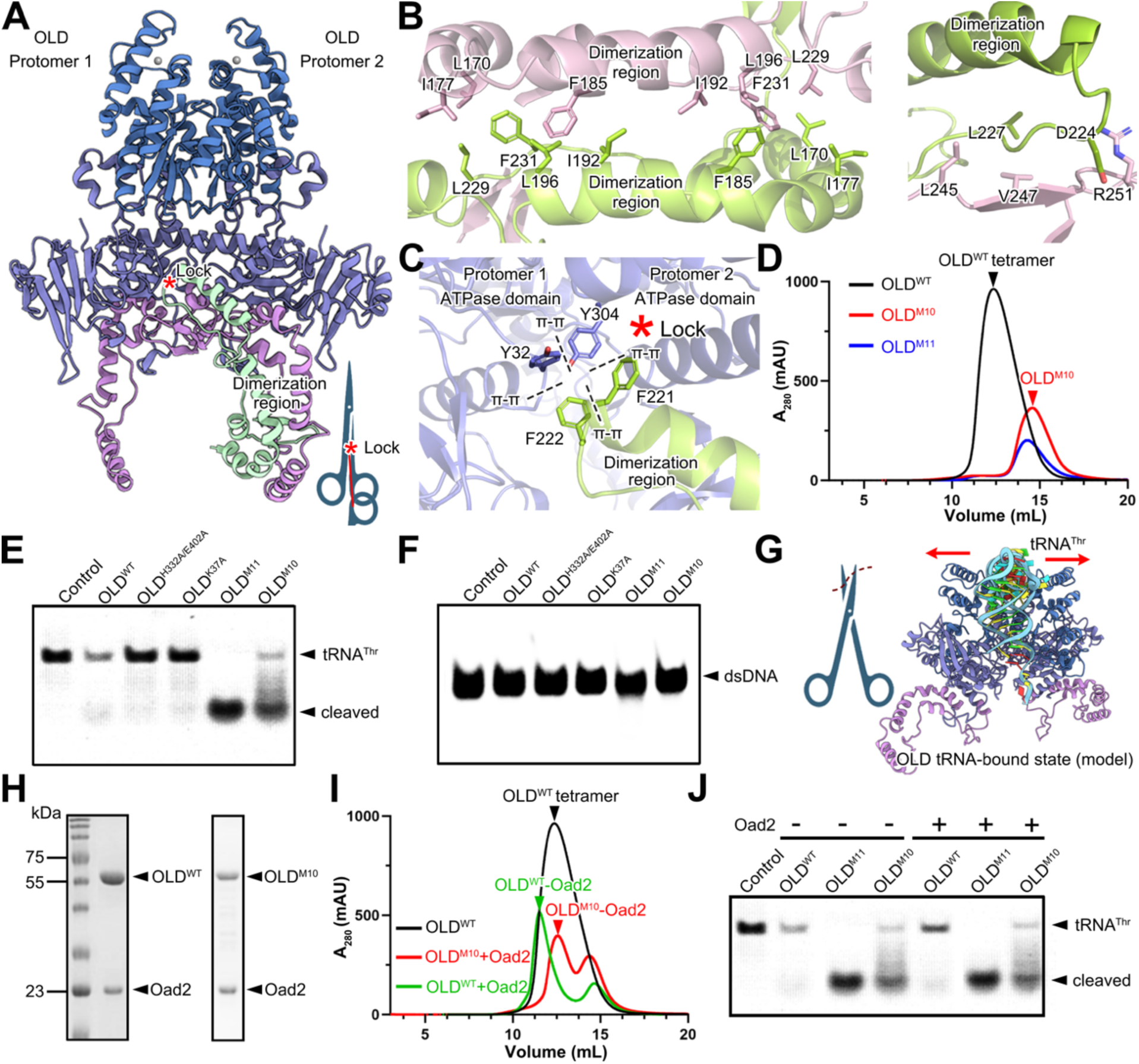
OLD dimer demonstrates enhanced tRNAse activity. **(A)** Structural basis of the “lock” mechanism within the OLD tetramer. For clarity, only two full-size protomers and the dimerization region of the third protomer are shown (domains colored as in Figure 3A). **(B)** Close-up views of the dimerization region interface detailing the symmetric interaction between the two protomers (pink and light green). **(C)** Close-up view of the lock interface showing interactions between the dimerization region (light green) and the ATPase domains of the Protomers 1 and 2 that constitutes the core of the “lock”. **(D)** Size-exclusion chromatography profiles of OLD^WT^, OLD^M10^, and OLD^M11^. **(E-F)** Nuclease activity assays of OLD^WT^ and mutants. (E) Cleavage of *in vitro* transcribed tRNA^Thr^. (F) Cleavage of linear dsDNA. **(G)** Schematic of an OLD protomer in an “open” active state bound to tRNA (substrate). **(H)** SDS-PAGE analysis of the pull-down experiment showing the interaction between OLD^WT^ or OLD^M10^ with the Oad2 protein. **(I)** Size-exclusion chromatography analysis of OLD^WT^, OLD^M10^-Oad2, and the OLD^WT^-Oad2 complexes. **(J)** tRNA cleavage assay comparing the activity of OLD^WT^, OLD^M10^, and OLD^M11^ in the absence or presence of Oad2.

Additionally, the dimerization region from one OLD dimer inserts into the interaction interface of the ATPase domain of the opposite dimer. This insertion is stabilized by residues F222 and F221 (located in the "tail" of the comma-like structure), which form π–π interactions with Y32 and Y304 of the opposing protomer, respectively, acting like a clasp that locks the OLD dimer interface (Figure 4C).

To assess the functional importance of these residues, we constructed two mutants: mutant M10 (L170D, I177D, F185A, I192D, L196D, and L229D), targeting residues mainly in the "body" of the comma-like region, and mutant M11 (M10 enhanced with additional substitutions Y32A, F221A, F222A, and Y304A), which includes residues both from the "body" and the "head". Gel-filtration experiments confirmed that both M10 and M11 shifted the oligomeric state of OLD from tetramer to lower molecular weight oligomers (Figure 4D). Although attempts were made to determine the cryo-EM structures of these mutants, technical challenges such as preferred orientation prevented high-resolution reconstruction.

We further compared the nuclease activity of the dimeric OLD mutants with that of the wild-type protein. The M10 mutant exhibited enhanced cleavage of the *in vitro* transcribed tRNA Thr(UGU) relative to the wild type, while M11 showed the strongest tRNA cleavage activity (Figure 4E). In contrast, mutations in key residues of the TOPRIM or ATPase domains completely abolished tRNA cleavage (Figure 4E). Moreover, expression of dimeric OLD mutants *in vivo* led to cell toxicity (Figure S6). Additionally, neither the wild-type nor dimeric OLD mutants displayed detectable dsDNA nuclease activity (Figure 4F), aligning with our earlier findings. These results suggest that the activation mechanism of OLD could be closely linked to alterations in its tetramerization interface. We further modeled the open conformation of OLD without the inhibition of dimerization region. Significant conformational changes were observed: not only did the TOPRIM domains undergo substantial rearrangement - increasing the cleft diameter from 13.5 Å to 19.6 Å (measured between E398 residues from two protomers) - but the ATPase domains also shifted accordingly (Figure 4G).

Interestingly, *in vitro* pull-down and gel-filtration assays demonstrated that the Oad2 inhibitor forms a stable complex with both wild-type OLD and its dimeric mutants, indicating a direct physical interaction that modulates OLD enzymatic activity (Figure 4H). Nevertheless, Oad2 effectively inhibited the activity of the OLD^WT^ tetramer but had little inhibitory effect on OLD^M^^10^ or OLD^M^^11^ dimers, implying that Oad2-mediated inhibition could interfere with activation of the OLD system (Figure 4J).

### OLD detects unresolved chromosome termini in RecBCD-deficient cells

OLD toxicity is activated by phage RecBCD inhibitors or in *ΔrecB* cells. To identify potential additional host factors influencing OLD activation, we performed a pooled CRISPR interference (CRISPRi) screen against all *E. coli* genes. The only genes depleted in the presence of OLD represented the RecBCD subunits (*recB*, *recC*, and *recD*) (Figure 5A). This specificity was confirmed by efficiency of transformation (EOT) assays, where a plasmid encoding OLD was not efficiently transformed into *ΔrecB*, *ΔrecC*, or *ΔrecD* strains, while transformation into wild-type or *ΔrecA* cells remained efficient (Figure S7A, S7B). These results suggest that OLD either directly monitors the RecBCD complex integrity or detects the consequences of RecBCD depletion, such as accumulation of damaged DNA and unprotected dsDNA ends.

**Figure 5.**
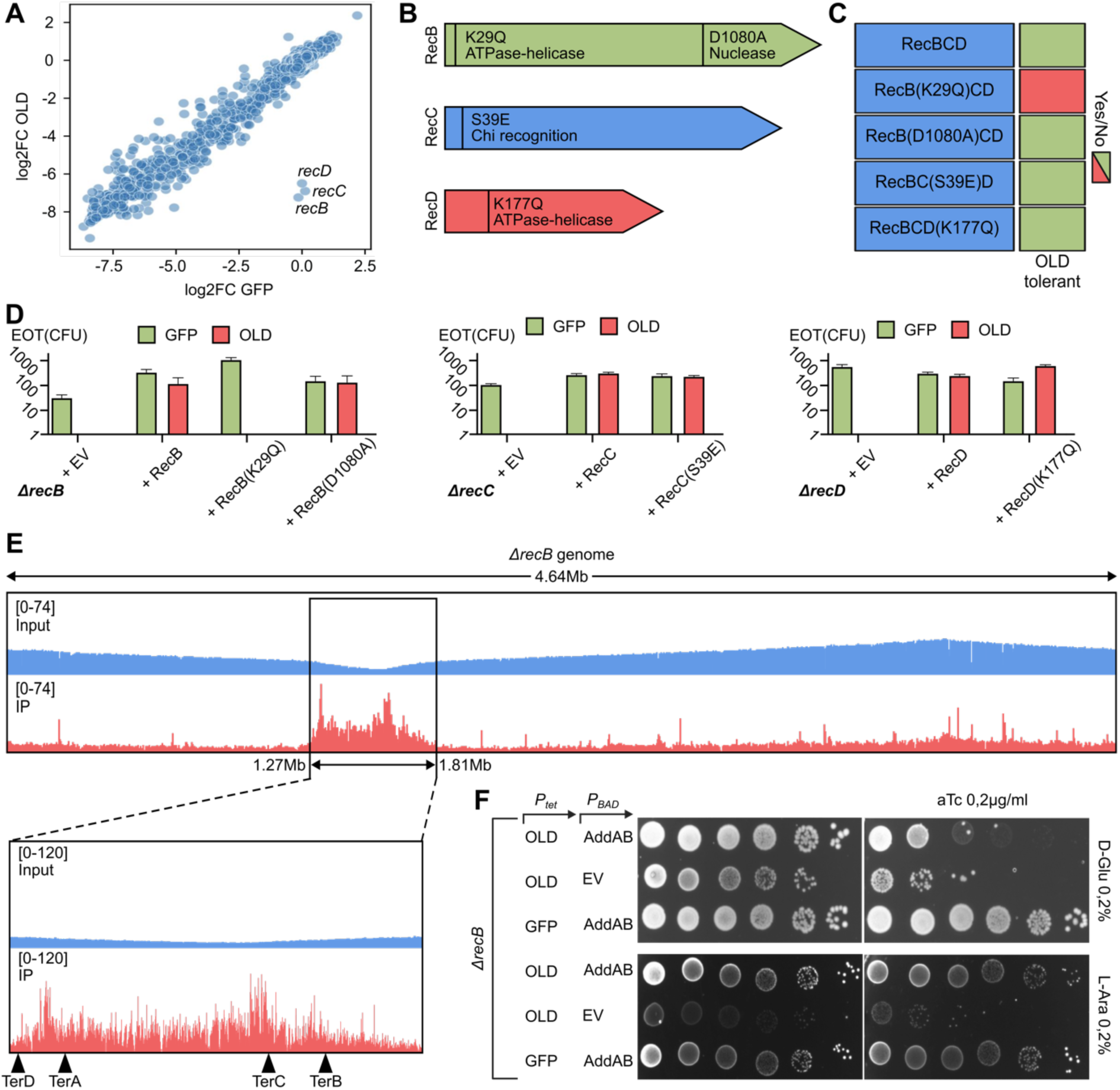
OLD surveils functionality of RecBCD complex by detecting aberrant DNA structures. **(A)** CRISPR interference (CRISPRi) screen identifies *recB*, *recC*, and *recD* as the only genes whose depletion is enriched in cells expressing OLD. **(B)** Scheme of the RecBCD complex and its catalytic mutants used in the efficiency of transformation (EOT) assays: nuclease-deficient RecB (D1080A), ATPase/helicase-deficient RecB(K29Q), χ-site recognition mutant RecC (S39E), and helicase-deficient RecD (K177Q). **(C, D)** EOT assay reveals that OLD toxicity is suppressed by wild-type RecBCD and all tested mutants except for translocation deficient RecB(K29Q). Controls without induction are demonstrated in Figure S7. **(E)** Chromatin immunoprecipitation sequencing (ChIP-seq) of OLD in Δ*recB* cells demonstrates specific binding to the chromosomal terminus region. **(F)** Viability of Δ*recB* cells expressing OLD is restored by heterologous expression of the *B. subtilis* AddAB complex.

To distinguish these models, we tested whether the physical presence or the enzymatic function of RecBCD was required to suppress OLD toxicity. First, we complemented strains with *recB*, *recC* or *recD* deletions with plasmids expressing wild-type RecBCD or specific mutants defective in nuclease activity [RecB(D1080A)]^88^; RecB helicase/ATPase activity [RecB(K29Q)]^89^; χ-site recognition [RecC(S39E)]^90^, or RecD helicase/ATPase activity [RecD(K177Q)]^91^(Figure 5B). The cells were transformed with OLD-carrying plasmid and CFU number was measured (Figures 5C, 5D, S7C and S7D). Only one mutant, RecB (K29Q), which is deficient in ATPase activity and translocation, failed to suppress OLD toxicity (Figures 5C, 5D, and S7D). This indicates that OLD does not monitor the mere presence of RecBCD but rather detects a loss of its function, suggesting that OLD senses the accumulation of unrepaired DNA structures when RecBCD translocation activity is impaired.

To identify the DNA substrates bound by OLD *in vivo*, we performed ChIP-seq with Flag-tagged OLD (E402A) in Δ*recB* cells. OLD binding sites were found to be enriched in the terminus region of *E. coli* chromosome (Figure 5E). This suggests that OLD detects aberrant DNA structures that accumulate at unresolved DNA termini in the absence of RecBCD. This finding could explain OLD dependency on RecD presence. While RecD-deficient cells could process dsDNA breaks and the RecBC complex retains translocation and recombination activity^92^, the primary defect of *ΔrecD* cells is the failure to properly resolve replication forks at the terminus and accumulation of over-replicated DNA^93^. Therefore, OLD toxicity is triggered whenever RecBCD function is compromised, irrespective of the specific mechanism, whether through the loss of DSB repair (Δ*recB/C*) or through impaired resolution of replication forks (Δ*recD*).

Consistent with this, heterologous expression of the AddAB complex, a functional RecBCD analog from *Bacillus subtilis*^94^, alleviated OLD toxicity in *ΔrecB* cells (Figure 5E). Furthermore, the *E. coli* JC5183 strain^95^, which lacks RecBC activity (*recB21, recC22)*, but carries the *sbcA5* mutation activating the alternative RecET pathway from prophage Rac, was fully compatible with OLD as shown by EOT (Figure S7E). Notably, OLD retained anti-phage activity in the RecBC^-^ background of JC5183, although it was lower than in the *E. coli* BW25113 host (Figure S7F). This suggests that while functional RecBCD activity is essential to prevent OLD activation in uninfected cells, OLD can detect phage infection not only through RecBCD inhibition, but potentially through alternative phage-specific DNA triggers.

### OLD senses phage replication via recognition of ssDNA hairpins in the λ phage origin

To further investigate how recombination systems contribute to OLD activation, we examined recombination-deficient mutants of λ in the OLD_Low_ background. In contrast to OLD expressed from a native promoter, OLD_Low_ did not completely restrict wild-type λ, while λ1188 (*gam^+^ red^-^*) was fully restricted (Figures 6A and S8A). This pattern indicates that the λ Red system protects the phage from OLD, as its absence sensitizes λ to restriction.

**Figure 6.**
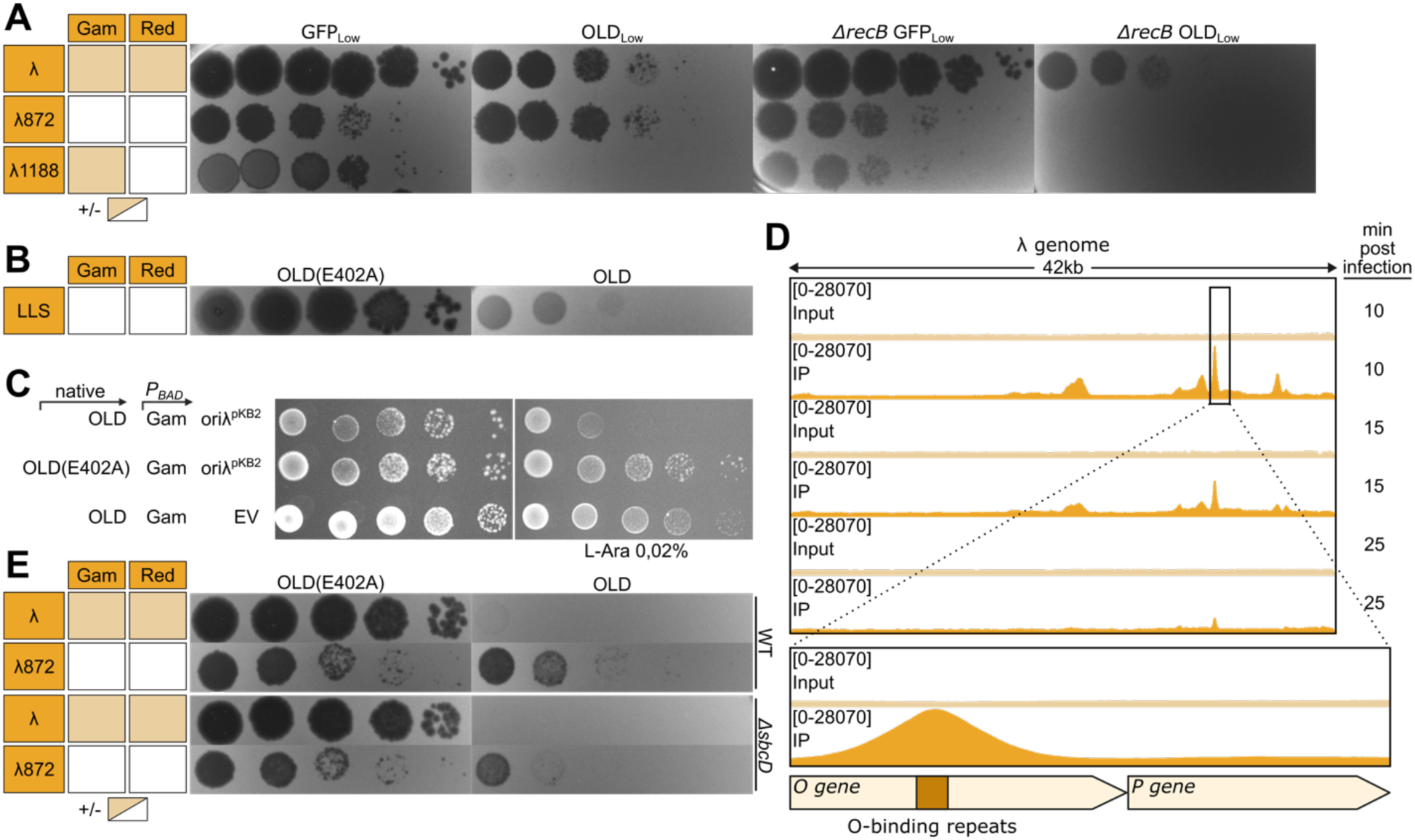
OLD is activated by ssDNA hairpins formed at the λ phage replication origin *in vivo*. **(A)** OLD-mediated restriction is modulated by phage-encoded recombination systems. Phage λ mutants lacking functional recombination machinery (*red⁻*) exhibit enhanced sensitivity to OLD, indicating that phage recombination proteins attenuate activation. **(B)** The LLS phage, which lacks its own RecBCD inhibitors and recombination system and relies on host RecBCD via 23 χ-sites, is restricted by OLD in a RecBCD⁺ background. This indicates that OLD activation during phage infection does not strictly require RecBCD inhibition. **(C)** OLD toxicity is potentiated by the presence of a plasmid carrying the λ replication origin (*oriλ*) and O/P replication proteins, demonstrating that the origin itself constitutes an activation signal. **(D)** Chromatin immunoprecipitation sequencing (ChIP-seq) during early λ infection reveals specific OLD binding to the O-binding repeats (iterons) within *oriλ* **(E)** OLD restricts the otherwise resistant λ872 (*gam^-^*) phage in a RecBCD⁺ host upon deletion of *sbcD*, which encodes a hairpin resolvase. This demonstrates that ssDNA hairpin accumulation is sufficient to trigger OLD activation even when RecBCD is fully functional.

Host recombination activity is required to resolve λ replication intermediates, if the phage does not encode λ Red system^26,96^, and we further tested phage growth in an OLD_Low_ *ΔrecB* background. Notably, the OLD_Low_ plasmid is tolerated by the *ΔrecB* cells. In these conditions, both λ1188 (*gam^+^ red^-^*) and λ872 (*gam^-^red^-^*) were completely restricted by OLD_Low_, while wild-type phage retained partial infectivity. Therefore, λ phage can productively infect OLD_Low_ cells if at least one recombination system (λ Red or RecBCD) is active. It can be surmised that recombination systems and OLD compete for the same DNA substrates, possibly dsDNA ends, generated during phage infection. When recombination is defective, these substrates accumulate and activate OLD.

The role of dsDNA ends as a possible substrate causing OLD activation is supported by isolation of escaper mutants of the T5-like Bas34 phage. While Bas34 demonstrates moderate sensitivity to OLD, resistant escapers acquired point mutations in homing nuclease *gp56* or *a1* genes (Figures S8C and S8D). A1 is an essential nuclease responsible for the host DNA degradation, therefore causing accumulation of free dsDNA ends^97^. Altogether this data suggests that phage recombination systems can resolve dsDNA-breaks and therefore act as anti-OLD systems.

However, expression of dsDNA-binding protein T4 gp2 does not inhibit OLD defense against λ (Figure S8B), and under native expression conditions OLD fully restricts wild-type λ (Figure S8A), despite the presence of a functional recombination pathway. Moreover, we tested the LLS phage – a lambdoid phage that fails to efficiently infect *ΔrecB* cells^98^. LLS lacks its own RecBCD inhibitors and recombination system, encodes 23 χ -sites and therefore relies entirely on host RecBCD for its replication. Strikingly, OLD was active against the LLS phage, despite the RecBCD^+^ background during infection (Figure 6B).

We hypothesized the existence of an additional, infection-specific OLD activation trigger, and attempted to isolate novel escapers of different phages. This resulted in isolation of the three OLD-resistant mutants of the phage HK578 (Figures 6B and S6C). One mutant (esc3) harbored a single nucleotide deletion in a *gp43* gene, encoded close to putative exonuclease and SSB protein, suggesting it’s involvement in recombination function. In contrast, two other escapers carried mutations related to replication initiation: esc1 contained a point mutation in the helicase-primase Gp46, and esc2 had a deletion in the promoter region of the operon encoding *gp45-gp46* genes. This pointed at phage replication as a possible source of DNA structures causing OLD activation.

Consistently, prior findings indicated that OLD toxicity during λ prophage induction required an intact replication origin (*oriλ*), viral O and P replication initiation proteins, and the host DnaB helicase^41^, but not DNA polymerase or phage genome replication. λ replication origin contains four palindromic sequences (iterons)^99^ that, upon dsDNA melting, can form ssDNA hairpin structures (Figures S9A and S9B). Activation by plasmid- or phage-specific replication *ori* ssDNA hairpins have been recently demonstrated for a Lamassu phage defense system^100,101^, also relying on ABC ATPase sensor domain. We tested whether activity of the phage replication *ori* contributes to OLD activation by co-expressing OLD with a theta-replicating plasmid (pKB2)^102^ carrying *oriλ* and the O/P proteins. While this plasmid alone was mildly toxic in OLD^+^ cells, even weak expression of Gam led to robust cell death, an effect not seen with a control plasmid (Figure 6C). This synergy indicates that an active phage origin dramatically potentiates OLD activation. To directly probe this interaction, we performed ChIP-seq of Flag-tagged OLD(E402A) during λ infection, as well as ChIP-seq with the strain carrying pKB2 *oriλ* plasmid. In both conditions, specific binding of OLD to the *oriλ* region was observed (Figures 6D and S9C). At the same time, no specific binding of OLD to the bacterial terminus regions during λ infection has been detected (Figure S9D). This observation suggests that despite RecBCD inhibition in the span of phage infection aberrant DNA structures in bacterial genome do not accumulate or not serve as a primary trigger for OLD.

SbcCD is a host enzyme that processes ssDNA hairpins. Gam protein was previously reported to inhibit SbcCD enzyme^103–105^ and thus Gam expression not only exposes dsDNA ends due to RecBCD inactivation but also stabilizes ssDNA hairpins through SbcCD inactivation. Because OLD binds specifically to the iteron-rich *oriλ*, we reasoned that SbcCD activity might suppress OLD activation by processing the hairpins, when OLD^+^ culture is infected with λ872 phage (lacking *gam*). Consistent with this, OLD-mediated defense was potentiated in a *ΔsbcD* host: the previously resistant λ872 phage became OLD-sensitive, despite the presence of active RecBCD. This result confirms ssDNA hairpin accumulation as an alternative and sufficient signal for OLD activation and explains how (*gam^-^ red^-^)* λ872 remains “invisible” for OLD in wild-type cells, but becomes sensitive in *ΔrecB* cells (due to exposed DNA ends) or in *ΔsbcD* cells (due to stability of ssDNA hairpins in the origin) (Figure 7).

**Figure 7.**
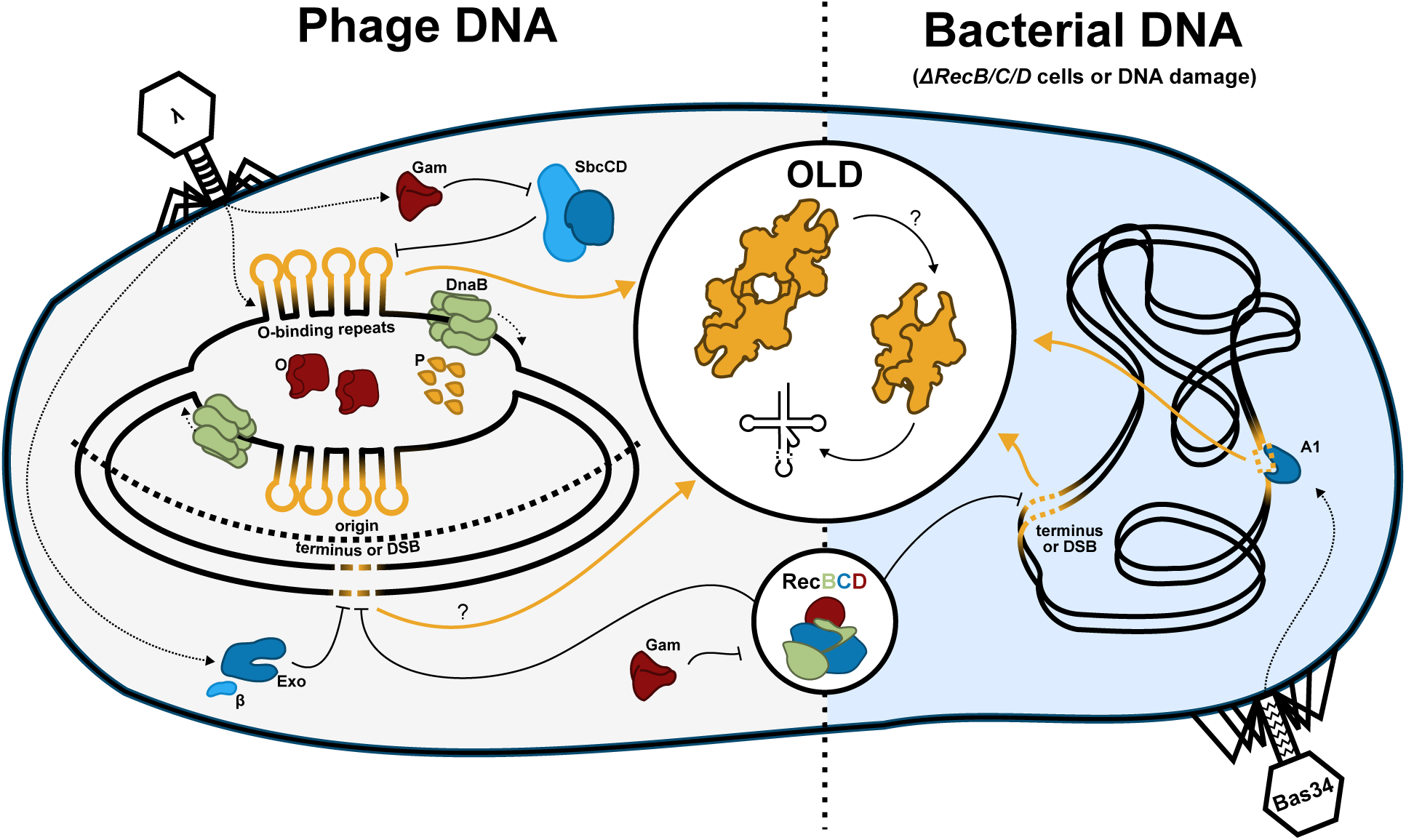
A unified model for OLD activation and function. OLD exists as an autoinhibited tetramer. Activation is triggered by various signals: ssDNA hairpins at the phage replication origin (*oriλ*), and unresolved DNA ends that accumulate in RecBCD-deficient cells, upon RecBCD inhibition, or due to phage nucleases such as A1 that generate dsDNA breaks. Both signals promote tetramer dissociation into active dimers, opening the TOPRIM catalytic site and triggering cleavage of host tRNAs. This arrests translation and aborts infection. SbcCD and phage/host recombination systems counteract activation by processing DNA substrates.

## Discussion

OLD of the prophage P2 is a prototypical member of the large family of proteins combining an ABC ATPase domain with an effector nuclease domain, most often TOPRIM^46^. This architecture is one of the most widespread in known and predicted immunity systems and is often encountered in proteins involved in metabolic functions, like DNA repair and replication control^46,63,72,74^. In this study, we determined the cryo-EM structure of the tetrameric OLD complex, demonstrating how ABC domains contribute to the control of the TOPRIM effector via oligomerization. In the systems encoding independent ABC and nuclease subunits, like PARIS and Lamassu, trigger sensing leads to the activation of the effector via its release from the inactivated complex and multimerization^18,19,70,71^. In contrast, the OLD-retron and Gabija systems, encoding fused ATPase and nuclease domains, regulate effector activation via conformational changes of the complex, controlled by ATP binding, that do not alter its oligomeric state^50,51,54–57^. P2 OLD could potentially represent a distinct activation paradigm, where activation could rely on the dissociation of the tetramer. Notably, a related OLD protein from *Thermus scotoductus* was reported to predominantly exist in a dimeric state^44^. While further experimental evidence is needed to verify this model, we demonstrate that mutations disrupting the OLD dimer:dimer interface result in catalytically-active protein and opening of the TOPRIM catalytic cleft, a transition likely intercepted by viral anti-OLD proteins, as evidenced by the inability of Oad2 to inhibit the *in vitro* activity of the OLD dimer.

Previous reports focused on OLD protein’s activity against DNA substrates^42–45^, however, in the early works about P2 OLD toxicity, translation inhibition and decrease in tRNA charging ability was noticed^39–41^, while phage and plasmid DNA isolated from OLD-activated cells was shown to remain intact^41^. Using microscopy, RNA sequencing, *in vivo* northern blots and *in vitro* cleavage assays we confirm that OLD inhibits translation and attacks an anti-codon stem loop of a subset of host tRNAs, an activity that also has been studied in detail in a parallel study^106^. Notably, OLD introduces two nicks, leading to the complete loss of the anticodon arm, and this activity could represent an adaptation to phage tRNA repair systems that efficiently ligate nicked tRNA, but could not repair tRNA with amputated anti-codon arm^106^. We demonstrate that expression of a single OLD-resistant tRNA Thr(UGU) could inhibit OLD defense against some phages, suggesting that tRNA cleavage is the primary cause of OLD toxicity.

Cleavage of the host tRNAs is a common strategy among bacterial (Cas12^107^, Cas13^108^, PARIS^18^, retron-Septu^109,110^, RemAIN^111^, HepS^112^, RAZR^113^, RloC^66^, ApeA^114^, DARNA^115^, Schlafen^116^) and eukaryotic (SAMD9^117^) anti-viral immunity systems, first described for the PrrC anti-codon nuclease^67^. In addition, TA modules with anti-phage activity, like MenT^118^ and CapRel^119,120^ can inactivate tRNAs via acceptor stem modification^121^. This mode of immunity can be subverted by phages via expression of tRNA variants resistant to cleavage^18,86,122^, or via tRNA fragments ligation^123,124^. Often a specific set of tRNAs is targeted by immunity systems, making it challenging for a phage to overcome the defense and requiring expansion of the tRNA gene set. We demonstrate that OLD preferentially targets Thr(UGU), Gly(UCC), fMet(CAU), Val(UAC), and Pro(UGG) tRNAs, while Thr(UGU) from T4, HK446 and T5-like phages inhibited OLD defense. At the same time, expression of a single tRNA was insufficient to alleviate OLD toxicity in *ΔrecB* cells suggesting that a full set of OLD-targeting tRNAs is required for complete OLD suppression.

Notably, while T5- and T4-like phages encode arrays of tRNA genes, HK446 encodes a single OLD-resistant Thr(UGU) and in addition carries an anti-OLD protein Oad2; therefore, its tRNA could rather represent an adaptation to the temperate lifestyle, since prophages often express tRNAs to compensate for the disrupted host tRNA genes used for integration^125^. Interestingly, among *Tequintavirus* phages Bas34 was the only restricted by OLD. Genome analysis revealed that in contrast to relatives, Bas34 lacks four out of five tRNAs targeted by OLD, including tRNA Thr(UGU), which we demonstrated to rescue OLD-sensitive phages (Figure S10). This highlights how phage-encoded tRNA arsenals shape resistance patterns towards tRNA-targeting immune systems.

We further explored the mechanisms of OLD activation and its connection with the host RecBCD function. OLD toxicity is triggered in *ΔrecB* cells or upon expression of known viral RecBCD inhibitors λ Gam and T7 Gp5.9. We identified a novel OLD trigger, Gp33 from the phages HK75 and HK446, and an OLD inhibitor Oad2, both encoded in the recombination-related genomic locus. In various “lambdoid” phages, this region is known for its high modularity^98,126^, and likely represents a hotspot mediating conflicts between host and viral recombination systems. As a second layer of complexity, immunity systems, like OLD and some retrons (Ec48^28^, Se72^29^, and Type VI^81^), can be activated in response to altered recombination activity of the host. This could drive the acquisition of inhibitors of such systems in viral recombination-related loci. Notably, Oad2 is related to the phage P22 EaE protein and in some phages can be fused with an HNH homing nuclease^127^ or an EaD/Ea22 protein^128^, previously studied in the context of λ lysogenization function^129,130^. Our results suggest that recombination-related hotspots could represent a rich reservoir of novel RecBCD inhibitors and quickly evolving anti-immunity proteins.

How exactly does OLD sense inhibition of RecBCD function? Our data indicate that this is an indirect process, i.e., that OLD is triggered by DNA substrates accumulating in the cell in the absence of RecBCD activity. RecBCD is a nuclease-helicase responsible for dsDNA breaks recognition, processing, and homologous recombination^22,131^. First, we demonstrated that the RecB(K29Q)CD translocation-deficient complex could not suppress OLD toxicity, ruling out the hypothesis of physical monitoring of RecBCD integrity. Second, we identified that the RecD subunit is required for suppression of OLD toxicity, despite the RecBC complex being active in DNA translocation and recombination^92^. Overreplication of the chromosome and accumulation of aberrant DNA structures at the terminus region is the most notable defect associated with *recD* deletion^93^. Lastly, we demonstrated that RecBCD deficiency could be complemented by alternative recombination systems, like AddAB and RecET. To confirm direct interaction with DNA substrates, we performed OLD ChIP-seq in *ΔrecB* cells, that demonstrated enrichment of OLD binding sites in the terminus regions. *E. coli.* Ter region is known for genome instability and also serves as a predominant source of DNA fragments for CRISPR adaptation^132^ and Argonaute guides^133^.

Importantly, we found out that OLD retains its anti-phage function in *ΔrecB* cells, suggesting that additional activation signals emerge in the cell during phage infection. By selecting escapers, we identified that phages could overcome OLD defense via mutations affecting replication initiation. Considering that melting of the phage λ replication origin was previously shown to be essential for triggering OLD toxicity^40,41^, we hypothesized that OLD could recognize ssDNA hairpins associated with iteron regions in the replication origins of multiple phages. In support of this, pKB2 vector, that encodes phage λ *ori* and replicates via theta mode^134^, triggers OLD toxicity, while direct interaction of OLD with the λ *ori* was shown via ChIP-seq both in the presence of pKB2 and during phage λ infection. Finally, we demonstrate that ssDNA processing nuclease SbcCD and phage λ Exo, β proteins counteract OLD activation.

Together, our data outline a complex landscape of the OLD immunity activation by DNA, allowing OLD to directly sense viral DNA replication, and identify phage and host proteins that modulate OLD activity by competing for DNA substrates. Considering that at least one dsDNA end sensing system (λ Exo, β or RecBCD) should be present to support λ infection in OLD⁺ cells and considering that phage nucleases trigger OLD, we hypothesize that in addition to ssDNA hairpins OLD can be activated by generation of dsDNA ends. Disruption of the host DNA integrity activates distinct immunity systems, like RloC^135^, Hachiman^136^, and retrons^29^. Activation by phage DNA was also recently proposed for the OLD-related Gabija system, where phage dsDNA end binding proteins were shown to license GajAB-mediated recognition of the DNA, by excluding RecBCD^52^. Given that OLD-like proteins represent one of the most abundant groups in bacterial anti-phage defense, these results highlight an important role of RecBCD in safeguarding bacterial cells from autoimmunity.

We propose a unified model in which OLD toxic tRNA cleavage activity is triggered after recognition of a DNA substrate (Figure 7). During phage infection, two types of triggers could emerge: replication-associated ssDNA hairpins at the *ori* regions and unresolved DNA termini generated in the phage or host genome. Available dsDNA ends could be rapidly processed by phage recombination proteins or the host RecBCD enzyme, which masks them from recognition by OLD, while SbcCD resolves ssDNA hairpins. It could be possible that exposed dsDNA ends also produce ssDNA hairpin structures upon resection, alternatively, OLD could retain an ability to bind both types of DNA substrates, considering that ssDNA hairpin ends structurally resemble dsDNA termini^137,138^. Importantly, during phage λ infection ssDNA hairpins are still favored. By being able to sense both types of substrates OLD integrates diverse signals associated with phage replication and aberrant DNA processing, therefore extending its phage recognition range. Further *in vitro* and structural studies are needed to validate this model and identify molecular details of OLD activation in response to DNA.

## Methods

### Bacterial strains, phages and plasmids

The phages, bacterial strains and plasmids used in the study are listed in Tables S2, S3, S4 and the primers are listed in Table S5. Infection with phages was carried out in Luria-Bertani (LB) medium supplemented with 5 mM CaCl2, 10 mM MgCl2, 0,2% maltose, if demanded for phage adsorption. Unless otherwise indicated, 0,2% L-ara, 0.1 mM isopropyl-ß-D-thiogalactopyranoside (IPTG) and 0.2 µg/ml anhydrotetracycline (aTc) were used for induction.

### Efficiency of plating (EOP) assay

The activity of OLD on phage defense was measured in comparison to its catalytically dead E402A mutant. *E. coli* BW25113 carrying a plasmid either with OLD cloned under its native promoter or with its catalytically dead mutant was grown overnight in 5ml LB + tetracycline (10 µg/ml). Bacterial lawns were prepared by mixing 100µl of a stationary culture with 10 ml of LB + 0,6% agar supplemented with 5 mM CaCl2, 10 mM MgCl2, 0,2% maltose, if demanded for phage adsorption, and the mixture was poured onto a Petri dish with a solid bottom LB + 1,2% agar layer. Tenfold serial dilutions of phage stock were spotted on each plate and incubated overnight at 37 C.

### Toxicity assays

OLD toxicity in the presence of viral proteins was measured using a spot-test assay. *E. coli* BW25113 carrying a plasmid either with OLD cloned under its native promoter or with its catalytically dead mutant carrying the pBAD vector with a trigger were pregrown in 5 ml of LB at 37 C to OD600 0.6. Cultures were then plated on minimal-medium (M9 with 7.5% LB) agar plates (if LB agar plates not mentioned) supplemented with 0.2% L-ara by serial tenfold dilution. Control plates without induction contained 0.2% glucose to prevent leakage of the araBAD promoter.

### Phage genomes alignments

Phage genome alignments were performed using Clinker genome analyses. The phage nucleotide sequences were uploaded to Clinker^139^ with minimum alignment sequence identity set to 0.22. Coloring was done manually.

### DNA Library Construction and OLD Inhibitor Screening

The DNA library for subsequent OLD inhibitor screening was constructed using genomic DNA of the HK446 bacteriophage, which was isolated by the standard phenol-chloroform extraction method. Genomic library was prepared by Mosaic Ends Tagmentation (METa) Assembly^140^. Shortly, this method is based on DNA fragmentation using the enzyme transposase Tn5, which cuts the targeted dsDNA and then flanks the DNA fragments with short oligonucleotides, which are then used to clone the resulting inserts into an expression vector.

We performed tagmentation using in-house-purified Sumo-tagged transposase enzyme, related to the hyperactive allele Tn5 containing classical mutations E54K and L372P. Purified Sumo-Tn5 stock contained approximately 750 ng/μl protein was loaded by annealed Mosaic end primers 5Phos_METagA1 and METagA2 (Evrogen) in concentration of 100 μM in 50 mM NaCl, 40 mM Tris-HCl (pH 8.0). Loaded transposomes were prepared by combining 0.141 volumes of annealed mosaic ends oligos with one volume of 750 ng/µl enzyme stock and incubating at room temperature (ca. 23°C) for 1 h. Tagmentation reaction was carried out with 10 μg purified gDNA of HK446 phage in volume of 1 ml by mixing MilliQ water, 5X TRIS-DMF buffer (50 mM TRIS-HCl buffer (pH 8.5), 25 mM MgCl2, 50% v/v dimethylformamide) and 130 μg of loaded transposome. Reactions were incubated for 7 min at 55°C at which point reactions were quenched by addition of SDS to the final concentration of 0.05% and incubation at 55°C for 5 min. The resulting mixture of DNA fragments was purified and concentrated on silica column with PCR & DNA Cleanup Kit (Monarch, NEB) and then separated in an 0.7% agarose gel, followed by cutting out a region of the gel in the range of 0.5-5 kb. The DNA fragments were then purified from the gel using DNA Gel Extraction Kit (Monarch, NEB). Next, eluted DNA inserts were subjected to an end-filling step by Q5 high-fidelity polymerase (NEB) with incubation at 72°C for 15 min to fill in 5’ overhangs. DNA fragments were then purified by silica column as before. Assembly reactions were prepared with the NEBuilder HiFi DNA Assembly enzyme mix following the manufacturer’s guidelines, using an approximate 2:1 molar ratio of the DNA insert to the vector. Cloning was carried out into the pFD-ME vector (CmR), linearized by PCR with primers pFD_Lin_F and pFD_Lin_R (Table S5), which are homologous to the flanking regions of the DNA inserts.

The assembly reaction in volume of 100 μl was desalted by PCR & DNA Cleanup Kit (Monarch, NEB), eluted with 25 μl 55°C water (the ∼1 μg assembly), and electroporated into 100 μl of NEB 10-beta electrocompetent *E. coli* cells at 1.8 kV. Cells were immediately rescued in 1ml of 37°C SOC outgrowth medium (0.5% Yeast Extract, 2% Tryptone, 10 mM NaCl, 2.5 mM KCl, 10 mM MgCl2, 10 mM MgSO4, 20 mM Glucose) and left on recovery with shaking at 37°C for 1 h. Next, 100-, 1000- and 10,000-fold dilutions of cells were plated onto LB (Cm 30 μg/ml) plates overnight at 37°C. The remaining DNA library culture was incubated overnight in 30 ml LB (Cm 30 μg/ml) at 18°C and, the next day, transferred to 37°C shaking incubator until it reaches an optical density at 600 nm (OD600) of between 0.6 and 1.0 AU. The amplified library was concentrated by centrifugation (4000 g, 7 min), resuspended in 10 ml of LB (Cm 30 μg/ml) with 15% glycerol and aliquoted 1ml into cryotubes for a long storage at -80°C. The quality of obtained library was assessed by counting the CFU number per 1 ml total recovery volume, as well as determining the average length of cloned fragments using colony PCR. For PCR, 10 colonies were selected, the reaction was carried out with 5x PCR screen mix (Evrogen) and primers pFD_check_F / pFD_check_R(Table S5). The library size was approximately 15 × 10⁶ CFU/ml, with an average insert size of 1.2 kb.

To identify anti-OLD-encoding DNA fragments from the HK446 phage library, a plasmid mixture was isolated from an overnight culture using a miniprep kit and electroporated into *E. coli* strain AB1157*ΔrecB*. The transformed library yielded 1,500,000 CFU. After pooling, the transformants were grown in liquid LB medium supplemented with the pFD vector inducer DAPG (100 μM) to OD600 of 0.8 AU. Electrocompetent cells were then prepared from these cultures, into which plasmids pBR_OLD (TetR) or pBR_OLD(E402A) (mut) (TetR), as a negative control, were transformed. The cells were subsequently plated on Petri dishes containing Tet (20 μg/ml), Cm (30 μg/ml), and DAPG (100 μM), and incubated overnight at 37°C. The next day, from the plates containing pBR_OLD co-transformed with the induced HK446 library, ten single colonies were selected for further screening of plasmids carrying potential anti-OLD genes. Candidate plasmids were purified using a plasmid miniprep kit (Evrogen) and transformed into *E. coli* strain BW25113 already harboring plasmids pBAD-Gam (AmpR) and pBR_OLD (TetR); the empty pFD vector served as a control. The resulting strains were assessed for toxicity after growth in liquid LB to OD600 = 0.3, using 10-fold serial dilutions plated on Petri dishes containing DAPG (100 μM) along with either arabinose (0.2%) to induce gam gene expression or glucose (0.2%) to repress it. pFD plasmid variants conferring protection in the presence of gam were selected and sequenced by Sanger sequencing.^105^

### Bacterial growth in liquid media

Growth of bacterial cultures to study phage infection or toxicity of SAMase was carried at 37 C in a 96-well format using EnSpire Multimode Plate Reader (Perkin Elmer). E. *coli* BW25113 carrying a plasmid either with OLD cloned under its native promoter or with its catalytically dead mutant were pregrown in 5 ml of LB supplemented with 10 mM MgCl2, 0,2% maltose at 37 C to OD600 0.6. 180µl of cells were then transferred to the 96-well and supplemented with phage MOI (0, 0.1 or 5) and brought to a final volume of 200 µl with LB supplemented with 10 mM MgCl2, 0,2% maltose. Optical density was monitored for 10h. All experiments were performed in three biological replicas.

### Microscopy

*E. coli* BW25113 carrying a plasmid either with OLD cloned under its native promoter or with its catalytically dead mutant carrying the pBAD vector with a trigger were pregrown in 5 ml of LB at 37 C to OD600 0.3. Cultures were then supplemented with 0,5% L-Ara and grown for 90 minutes. An aliquot was mixed with DAPI (Invitrogen) at 1 mg/ml final concentration and incubated at room temperature in the dark for 5 min. LB + 1.5% agarose slabs (approximately 0.2 mm thick) were prepared on a 75 × 25 mm^2^ microscopy slide (Fisher Scientific). Agarose slabs contained staining dyes at the same concentration were optionally supplemented with 0.5% L-ara, to induce Gam expression. Roughly 1 μl of stained cells was placed on an agarose slab and, following 1 min of drying, the slab was covered by a small (22 × 22 mm^2^) coverslip (Fisher Scientific). Imaging was performed on a Nikon Eclipse Ti-E inverted epifluorescence microscope. Each field of view was imaged in the transmitted light channel (200 ms exposure) and in the DAPI (200 ms exposure) channels.

### Total DNA extraction

*E. coli* BW25113 carrying a plasmid either with OLD or with its catalytically dead mutant (E402A) carrying the pBAD vector with Gam were grown in 5 ml of LB at 37 C to OD600 0.3. Cells were then pelleted and resuspended in M9 minimal media containing 7,5% LB and 0,2% arabinose. Cells were grown at 37C for 2 more hours. 4ml of each cell culture was pelleted and mixed with 500µl TE buffer, 50µl 10% SDS and 5µl of proteinase K (ThermoFisher 20mg/ml). The solution was then incubated at 55C for 20 minutes. Then 5µl of RNase A (ThermoFisher 10mg/ml) was added and the solution was incubated for 10 minutes at 37C, while shaking. To the resulting solution 800µl of phenol-chloroform was added, gently mixed and then centrifuged at 8000g for 3 minutes. The phenol fraction was collected and 500µl of chloroform was added, the solution was gently mixed and centrifuged at 8000g for 3 minutes. The previous step was repeated 2 more times. Finally, to the collected phenol fraction 50µl of 5M NaCl, 2µl of glycogen and 1,3ml of cold EtOH(96%) was added. The probes were then incubated at -20C for 2 hours. After 2 hours the probes were centrifuged, the pellet was washed once in 70% ethanol, then dried and resuspended in 200µl of TE buffer. Equal amounts of DNA were then visualized on 0,5% agarose 1x TAE gel stained by ethidium bromide.

### Total RNA extraction

*E. coli* BW25113 carrying a plasmid either with OLD or with its catalytically dead mutant (E402A) carrying the pBAD vector with Gam were grown in the same conditions as described in DNA extraction method. Total RNA was purified using Evrogen RNA Solo Kit according to the manufacturer’s instructions. Equal amounts of RNA were then visualized on 1% agarose 1x TAE gel stained by ethidium bromide.

### Phage gDNA extraction

Phages was produced from single plaque in 10ml LB. The lysate was then incubated with 2µl of DNase I (ThermoFisher 50U/µl), 2µl RNase A (ThermoFisher 10mg/ml) for 1 hour at 37C, while shaking. Then 2ml 5M NaCl and 800mg PEG 8000 was added and incubated at 4C overnight with continuous linear shaking. The probes were then centrifuged at 4000g for 1 hour, supernatant was discarded. The pellet was then resuspended in 400µl STM buffer. Then 400µl of chloroform was added to elute PEG. The probes were vortexed for 2mins and then centrifuged at 10000g for 5 minutes. The supernatant was collected and treated with 16µl 0,5M EDTA, 4µl Proteinase K (ThermoFisher 20mg/ml) and 20µl 10% SDS for 90 minutes at 50C. Then 400µl of phenol-chloroform was added and gently mixed. The probes were centrifuged at 14000g for 3 minutes, phenol fraction was collected and repeatedly washed with phenol-chloroform solution for 2 more times. Finally, phenol fraction was supplemented with 40µl 3M sodium acetate (CH_3_COONa), 2µl glycogen and 1ml of ice-cold (96%). The probes were left overnight at -20C. The probes were then centrifuged, the pellet was washed once in 70% ethanol, then dried and resuspended in 100µl of TE buffer.

### TUNEL assay

*E. coli* BW25113 carrying a plasmid either with OLD or with its catalytically dead mutant carrying the pBAD vector with Gam were pregrown in 5 ml of LB at 37 C to OD600 0.3. Cells were then pelleted and resuspended in M9 minimal media containing 7,5% LB and 0,2% arabinose. Cells were grown at 37C for 2 more hours. Fixed cells were stained with terminal deoxynucleotidyl transferase dUTP nick end labeling (TUNEL) assay kit. Flow cytometry data were acquired on a Beckman coulter Navios and analyzed using FlowJo v10. Prior to analysis, a gate was applied to a FSC-H vs FSC-A plot to exclude doublets and debris. 200 000 events were collected for each sample in three biological replicates. A positive signal threshold for FITC was established based on the BW25113 as negative control, defined as the level exceeding the signal of 4.95% of cells in the control population. The frequency of TUNEL-positive cells in experimental samples was determined using this gating strategy and compared to the negative control.

### Preparation of enriched small-RNA fractions

Small-RNA–enriched fractions were prepared from cultures grown in the OLD system following a protocol adapted from our previous workflow. *E. coli* MG1655 carrying either pFR_OLD (Ptet–OLD) or the control plasmid pFR66 (Ptet–sfGFP) was used; strains also contained pFD265 (PhlF–gam), placing Gam under DAPG-inducible control.

Cells were diluted 1:100 in LB with the appropriate antibiotics from overnight cultures and grown with agitation at 37 C. When cultures reached OD600 0.2, aTc was added to a final concentration of 0.5 μg ml−1 to induce expression of the OLD system. Growth continued to OD600 0.4, and the gam trigger was induced with DAPG added to a final concentration of 50 μM, followed by incubation for an additional 30 min post induction. The pellets were suspended in 2 ml of 50 mM sodium acetate (NaOAc) and 10 mM MgOAc pH 5.0, with the addition of 1.9 ml of commercial acidic phenol pH 4.5 (Ambion). The resulting mixture was shaken at 200 rpm for 30 min at 37 C then centrifuged at 5,000g for 15 min at 4 C. The upper aqueous phases were collected and subjected to precipitation of total nucleic acids in 0.1 M NaCl, supplemented with an equal volume of 2-propanol, for overnight incubation at room temperature. Following centrifugation at 14,500g for 15 min at 4 C, pellets were washed with 80% ethanol and air-dried. Removal of rRNA from samples was achieved through precipitation: pellets were suspended in 0.8 ml of cold 1 M NaCl and spun at 9,500g for 20 min at 4 C. Supernatants were collected, mixed with 1.7 ml of ethanol, incubated for 30 min at −20 C and then centrifuged at 14,500g for 5 min at 4 C. The pellets were then washed with 80% ethanol and air-dried.

Removal of DNA from samples was achieved through precipitation: pellets were resuspended in 0.6 ml of 0.3 M NaOAc pH 5.0. The mixture was then heated for 5 min at 60 C with regular pipetting, followed by the addition of 0.34 ml of 2-propanol and incubation for 10 min at room temperature; the solution was then centrifuged at 14,500g for 5 min at room temperature and supernatants collected. The resulting small RNA fractions were precipitated by the addition of 0.23 ml of ethanol to the supernatant, centrifuging at 14,500g for 15 min at 4 C, washing the pellet with 80% ethanol and air-drying for 5–10 min. Pellets were dissolved in 0.35 ml of RNase-free water.

For deacylation, the fraction was incubated in 0.1 M Tris pH 9.0 for 45 min at 37 C. Final precipitation in 0.3 M NaOAc pH 5.0 was achieved by the addition of 2.7 volumes of ethanol, incubation for 30 min at −80 C and centrifugation at 16,000g for 25 min at 4 C. Pellets were washed with 80% ethanol, air-dried and dissolved in 80 μl of 1 mM sodium citrate.

### Small-RNA library preparation and sequencing

RNA was dephosphorylated with Quick CIP (NEB) at 37 C for 15 min and purified using the Monarch Spin RNA Cleanup Kit (eluted in 20 µL). 200 ng of dephosphorylated RNA was ligated to a pre-adenylated 3ʹ adapter (SB104) using T4 RNA Ligase 2, truncated KQ, in 1× ligase buffer with 14% PEG 8000, 16 pmol adapter, for 2 h at 25 C, followed by cleanup (Monarch kit, 20 µL elution). Reverse transcription was performed with 180 ng ligated RNA using Induro RT (NEB) and 20 µM (16 pmol) primer SB103 for 16 h at 37 C. The reaction was treated with 2U Exonuclease I for 30 min at 37 C, heat-inactivated at 80 C for 20 min, and subsequently treated with 5U RNase H and 10 µg RNase A for 20 min at 37 C.

Resulting cDNA was ligated to a 5ʹ adapter (SB077) using thermophilic 5ʹApp DNA/RNA ligase (NEB) in 1× ligase buffer with 5 µM adapter, overnight (16 h) at 65 C, followed by cleanup. Residual primers were degraded by sequential treatment with RppH (5U, 37 C, 30 min), Lambda Exonuclease (5U, 37 C, 30 min), and Exonuclease I (20U, 37 C, 30 min). Final cDNA was purified using the Monarch kit with 0.8× ethanol and eluted in 20 µL.

Libraries were amplified from 10% of the cDNA using indexed Illumina primers and Q5 DNA polymerase in 25 µL reactions for 25 cycles (60 C annealing, 30 s extension). Amplified libraries were size-selected using Beckman SPRI beads with a double-sided selection (0.7×/1.2× bead-to-sample ratios) to enrich fragments >85 bp and <200 bp. Libraries were quantified (TapeStation) and multiplexed for sequencing on an Illumina MiSeq i100 platform (paired-end, 2×150 nt reads). Reads were aligned to a curated *E. coli* tRNA and rRNA reference using Bowtie2 (--very-sensitive-local, retaining uniquely mapped reads only).

### tRNA-seq processing and analysis

The scatter plot was generated by parsing aligned reads from the Bowtie2-derived count table. For each tRNA in each replicate, read length was determined from genomic coordinates, and counts were summed to obtain total reads and long reads (>65 nt). For each tRNA, the proportion of long reads was calculated as the number of reads >65 nt divided by the total reads in the sample. These proportions were then compared between conditions using a log–log scatter plot. tRNAs showing fold enrichment or depletion greater than twofold were highlighted in red, while all others were shown in grey.

The heatmap was generated using a custom pipeline. 5ʹ- and 3ʹ-end counts were aggregated for each tRNA species at each nucleotide position. Positional read counts were normalized using “3ʹ-CAA anchoring,” in which each tRNA’s profile is scaled by the number of reads terminating at its annotated mature 3ʹ-CAA end. This provides an internal reference that reduces biases arising from reverse-transcriptase drop-off at modified nucleotides or secondary structures. Differential matrices (OLD − GFP) for 5ʹ starts and 3ʹ ends were then computed after aligning all tRNAs to a shared index and positional coordinate set. To emphasize fold-like differences while retaining sign, the normalized values were transformed using: signed_log(x)=log10(500x+10)−log10(−500x+10)

The scatter plot was made by Aligned reads were parsed from the Bowtie2-derived count table,. For each tRNA and replicate, read length was computed from genomic coordinates, and counts were summed to obtain total reads and long reads (>65 nt). For each tRNA, the proportion of long reads was calculated as (>65 nt / sample read) and compared between conditions in a log–log scatter plot. FOLD enrichment/depletion >2 were highlighted by red dots, all others shown in grey.

### Northern blot

For Northern blot analysis, *E. coli* strains were grown in triplicate cultures inoculated from single colonies. After reaching OD 0,3 of cell culture samples were supplemented with 0,2μg/ml aTc and grown for another 40min. Then rapidly chilled on ice and harvested by centrifugation (8,000 to 10,000 x g, 4C). Total RNA was isolated using the TRIzol reagent (Invitrogen) according to manufacturer procedure. Precipitated RNA pellets were dried on air and were dissolved in nuclease-free water (Invitrogen). RNA concentrations were determined using a NanoDrop 2000c (Thermo Scientific) spectrophotometer. Prior to loading on a gel, the RNA (15 μg per sample for Northern blotting and 1 μg per sample for SYBR staining) was mixed with denaturing gel loading buffer (f.c. 1xTBE, 33% (*v/v*) formamide, 3 M Urea, 0.01% (*w/v*) bromophenol blue, 0.01% (*w/v*) xylene cyanol blue) followed by heating up to 95°C for 3 min and then immediate cooling on ice. After RNA separation by 12.5% PAGE (7 M urea), the gel was cut in two halves: the first one with 1 μg of total RNA samples was subjected to staining procedure (1x ssGreen RNA Gel Staining Solution (Lumiprobe) in 1xTBE, 5 min), the second one with 15 μg of total RNA samples was pre-incubated for 10 min in 0.5xTBE followed by the transfer on positively charged nylon membrane (Roche) by semi-dry electroblotting (Trans-Blot SD Cell, Bio-Rad) for 1h at 400 mA (0.5xTBE). For RNA immobilization the membrane was baking for 1 h at 80°C, then briefly rinsed in water and pre-incubated in 5 mL of hybridization buffer (DIG Easy Hybridization Granules, Roche) for 1 h at 42°C in a hybridization oven (AnalytikJena). The buffer was further replaced by a 5 mL of a fresh aliquot containing 2 nmol of fluorescently-labeled synthetic probe followed by overnight incubation at 42°C with continuous stirring. Hybridized membranes were washed 2х 5 min with washing buffer (2x SSC, 0.01% (w/v) SDS), briefly rinsed in water followed by detection of fluorescent signals (ChemiDoc MP Imaging System, Bio-Rad). Wet membranes were further pre-incubated with hybridization buffer and subjected for hybridization with second/third probe as described above. In case of hybridization with the next probe harboring the same fluorescent label the membrane was firstly stripped for 30 min at 98°C in a water bath in a stripping buffer (2x SSC, 1% (w/v) SDS), followed by brief incubation in water and detection of negative fluorescent background.

### CRISPRi

The CRISPRi library used to target the MG1655 genome was the EcoWG1 library^141^ which contains approximately 20,000 sgRNAs (five guides per gene). This EcoWG1 library was cloned into the pFR56 plasmid which carries dCas9 under the control of the 2,4-diacetylphloroglucinol (DAPG)-inducible pPhlF promoter and an RP4 origin of transfer to enable conjugation. This plasmid also contains a constitutively expressed sgRNA with two BsaI sites for library guides cloning via Golden Gate assembly. The plasmid was introduced by electroporation into *E. coli* MFDpir cells carrying pFR58 a low-copy pSC101, kanamycin-resistant plasmid expressing the PhlF repressor constitutively. This repressor prevents leaky dCas9 expression that could introduce a bias in the library.

MFDpir donor cells carrying both pFR56 (EcoWG1) and pFR58 were used to transfer the EcoWG1 library into recipient strains MG1655/pFR66 (sfGFP) or MG1655/pFR_OLD (Table S4). MFDpir donor cultures were grown to an OD ≈ 1 directly from 1 ml DMSO stock in 25 ml LB medium containing 0,3 mM diaminopimelic acid (DAP), 50 µg/m Kan and 20 µg/ml Cm. Plasmid DNA was extracted using the NucleoSpin plasmid kit (Macherey–Nagel) and used as a reference for the library composition before conjugation. Recipient strains were grown to stationary phase in LB with 50 µg/ml Kan. Donor and recipient cultures (1 ml of each) were centrifuged, washed (donor with LB+ DAP; recipients with LB) to remove antibiotics and mixed in a 1:1 volume ratio (0.5 ml each). The mixture was centrifuged (2,000g for 10 min), resuspended in 50 µl LB+DAP, spotted on LB+DAP agar plates, and incubated for 1.5 h at 37◦C. After conjugation, cells were scraped with an inoculating loop, resuspended in 1 ml of LB containing 20 µg/ml Cm plus 50 µg/ml Kan and plated on two 12cm× 12cm LB plates with the same antibiotics. Plates were incubated overnight at room temperature, and an aliquot was serially diluted and spotted in parallel to estimate conjugation efficiency. More than 10^7^ transconjugants were obtained for all strains. Transconjugants were resuspended in 5 ml LB and stored at −80◦C with 10% DMSO until screening.

From frozen stock, one hundred microliters of transconjugants with the library (MG1655/pFR66 (sfGFP) plus pFR56 (EcoWG1) or MG1655/pFR_OLD plus pFR56 (EcoWG1)) was inoculated into 5 ml LB + Cm + Kan in duplicate for each strain, and incubated overnight at 37°C with shaking. Plasmid DNA was extracted to serve as the reference samples for each strain. Three successive passages were then performed in LB medium containing 50 µM DAPG, each initiated by 1:100 dilution (50 µl into 5 ml), corresponding to approximately 20 generations. The cultures were incubated for 4 h after the first and second dilutions, and for 16 h after the last dilution. Plasmid DNA was again extracted from these cultures to obtain the induced samples with the final extraction at 16 h representing the time point.

Library sequencing was performed as previously described, with a few modifications. Briefly, primers listed in Table S5 were used to perform two consecutive PCR reactions with Q5 polymerase (New England Biolabs). Starting from 100 ng of library plasmid DNA, the first PCR (98°C for 3min; two cycles of 98◦C for 30 s, 60◦C for 90 s and 72◦C for 30 s; 72◦C for 5 min) was performed in 20 µl reaction with 2 µM of each primer (LC863 and primers PCR1-forward, see Table S5). PCR products were treated with 1 µl ExoI (New England Biolabs) during 1 h at 37°C to remove residual primers and purified with 20 µl AMpure XP magnetic beads (Beckman Coulter). A second PCR for Illumina library preparation was performed in a 30 µl using 1 µmol of primers LC415 and LC1516 (Table S5) with cycling conditions as follows: 95◦C for 3 min; 16× 98◦C for 10s, 66◦C for 30s and 72◦C for 30s; 72◦C for 5 min) to add the second index and the Illumina P5 and P7 adapters. The resulting 354-bp fragment of each sample was normalized to 30 ng per sample, pooled and gel purified. The final pooled library was sequenced on a Illumina NextSeq 500 sequencer using a custom protocol as described previously^142^.

### Efficiency of transformation (EOT) assay

Competent cells were produced according to standard Inoue protocol. Then the cells were transformed with a plasmid either with OLD cloned under its native promoter or with its catalytically dead mutant. Following transformation, cells were resuspended in LB broth to a final volume of 1 mL and allowed to recover for 1 hour at 37C with shaking. The cells were then pelleted and plated onto selective agar plates. Colony-forming units (CFUs) were quantified after overnight incubation at 37C to determine the transformation efficiency. For experiments using the inducible pFR_OLD plasmid, the 1 mL transformed culture was split into two equal aliquots after the recovery period. These were pelleted and plated onto two types of selective plates: one containing anhydrotetracycline (aTc), and one without aTc.

### Isolation and sequencing of escaper mutants

*E. coli* BW25113 carrying a plasmid with OLD cloned under its native promoter were grown in LB supplemented with 10mM MgCl_2_, 5mM CaCl_2_ till OD600 reached 0.6, cells were infected with HK578 or Bas34 at an approximate MOI of 0.1 and the culture incubated overnight at 37 C. Phage was collected the following day and 1 ml of lysate was used to initiate the next round of infection with fresh OLD culture. Escaper mutants were purified from single plaques obtained from OLD culture and produced on the *E. coli* BW25113. Ten ml of high-titer lysate (∼ 10^9^ plaque-forming units per ml) was used for phage genomic DNA purification as described previously.

Libraries for WGS sequencing were prepared from 500 ng of phage DNA using MGI Easy PCR-Free Library Prep Set (MGI Tech), following the manufacturer’s instructions. Enzymatic fragmentation was performed according to the manufacturer’s protocol, followed by selection of 400–450 bp-long fragments on the provided DNA Clean Beads. The concentration of the prepared libraries was measured using Qubit Flex (Life Technologies) with the dsDNA HS Assay Kit. The quality of the prepared libraries was assessed using 4200 TapeStation System with the High Sensitivity D1000 ScreenTape Assay (Agilent). DNA libraries were further circularized, pooled and sequenced using DNBSEQ-G400 in 2x150bp PE mode.

De novo genome assembly was performed with the Shovill pipeline (v1.1.0; Seemann, https://github.com/tseemann/shovill), which uses SPAdes^143^ at its core with optimized pre- and post-processing of the paired-end reads. Single-nucleotide polymorphisms (SNPs) relative to the reference genome were identified with Snippy (v4.6.0; Seemann, https://github.com/tseemann/snippy), which maps reads with BWA-MEM^144^, processes alignments with SAMtools^145^, calls variants with FreeBayes, and annotates their functional effects with SnpEff^146^.

### Protein purification

OLD protein was purified from BL21(DE3) cells carrying pRSF_OLD. 4 L cell culture was grown at 37°C in LB medium supplemented with kanamycin to OD_600_ = 0.8, induced with 1 mM IPTG at 16℃ for 20 hours, and harvested. Cells were resuspended in lysis buffer (20 mM Tris-HCl, pH 7.6, 200 mM NaCl, 5%(v/v) Glycerol) and lysed by sonication on ice. Cell lysate was cleared by centrifugation for 30 min at 17,000 rpm at 4°C, and applied to a 5 mL Ni-NTA column equilibrated with lysis buffer. Ni-NTA column was subsequently washed with lysis buffer containing 25 mM imidazole, and OLD protein was eluted with lysis buffer containing 200 mM imidazole. The eluted fractions was loaded into a 5 mL HiTrap Q HP column (Cytiva) equilibrated with QA buffer (20 mM Tris-HCl, pH 7.6, 5% (v/v) Glycerol, 1 mM EDTA, 1 mM DTT, 50 mM NaCl). OLD protein was eluted by a 0.05-0.5 M NaCl gradient over 20 column volumes, concentrated, and diluted with QA buffer to adjust NaCl concentration to 0.1 M. OLD protein was next applied to a 5 mL HiTrap HP Heparin column (Cytiva) equilibrated with QA buffer. OLD protein was eluted by a 0.05-0.7 M NaCl gradient over 20 column volumes and concentrated to 2 mL volume. Finally, OLD protein was applied to size-exclusion column (Superdex 200 Increase 10/300 GL, Cytiva) equilibrated with 10 mM Tris-HCl pH= 7.6, 200 mM KCl. Fractions containing pure OLD protein were combined and concentrated using 15 mL Amicon Ultra 50 kDa MWCO concentrator. The product (purity >98%) was stored in aliquots at -80 C. OLD protein derivatives were prepared by the same procedure.

Oad2 was purified from BL21(DE3) cells transformed with pET22b_Oad2. The protein expression was induced with 1 mM IPTG at 18 °C for 20 hours at OD_600_ of 0.7. The cell pellet was lysed in lysis buffer (20 mM Tris-HCl, pH 7.6, 200 mM NaCl, 5%(v/v) Glycerol) using sonication. The supernatant was loaded on a 5 mL Ni-NTA column that was subsequently washed and eluted with lysis buffer containing 250 mM imidazole. The eluted fractions were concentrated to 3 mL and applied to size-exclusion column (Hi-Load 16/600 Superdex 200 prep grade, Cytiva) equilibrated with 10 mM Tris-HCl pH 7.6, 200 mM KCl. The fractions containing target proteins were concentrated to 22 mg/mL, and stored at −80 C.

### Cryo-EM sample preparation

For cryo-EM grid preparation, OLD protein was concentrated to 14 mg/mL and mixed with CHAPSO (Hampton Research, Inc.; final concentration 8 mM) prior to grid preparation. About 3 μL samples were applied to freshly glow-discharged Quantifoil R1.2/1.3 holey carbon grids. The grids were blotted for 4.5 s with blot force 6 in a Thermo Fisher Scientific Vitrobot Mark IV and plunge-frozen in liquid ethane at liquid nitrogen temperature. The grids were negatively glow-charged using an EasyGlow Discharge System (PELCO) at 25 mA for 50s at 0.39 mBar. The ø 55/20 mm blotting paper used for plunge freezing was made by TED PELLA.

### Cryo-EM structure determination: data collection and data reduction

Cryo-EM data were collected at the cryo-EM core Facility of Shanghai Institute of Materia Medica (CAS), using a 300 kV Titan Krios (FEI/ThermoFisher) electron microscope equipped with a post-GIF Gatan K3 direct electron detector (Gatan). Data were collected automatically in the super-resolution mode, using EPU with a nominal magnification of 105,000x, a calibrated pixel size of 0.824 Å/pixel on the image plane, and with defocus values ranging from −0.8 to −2.0 μm. Each micrograph stack was dose-fractionated to 40 frames with a total electron dose of ∼50 e^−^/Å^2^ and a total exposure time of 2.1 s. 13661 micrographs were collected for further processing.

Data were processed as summarized in Supplementary Fig. 4. Data processing was performed using a Tensor TS4 Linux GPU workstation with four GTX 3090 graphic cards (NVIDIA). Dose weighting motion correction (5x5 tiles; b-factor = 150) were performed using Motioncor2 in Relion (4.0.1)^147^. Subsequent image processing was performed using Cryosparc (V4.6.2)^148^. Contrast-transfer-function (CTF) estimation was performed using Patch CTF Estimation in Cryosparc^149^. Automatic particle picking with Blob Picker and Template Picker yielded an initial set of 3,304,398 particles. Particles were extracted into 256 pixel boxes and subjected to rounds of reference-free 2D classification and removal of poorly populated classes, yielding a selected set of 379,579 particles. The selected set was kept for Ab-Initio reconstruction and further heterogeneous refinement with C2 symmetry. One among the four classes including 119,593 particles were selected for iterative rounds of homogeneous refinement and heterogeneous refinement with D2 symmetry, resulting in a 3.1 Å density map. The local resolutions of final maps were calculated by Local Resolution Estimation. Statistics for data collection and processing were summarized in Table S1.

The structure model for OLD tetramer was first predicted by Alphafold 2. Then initial model of OLD tetramer was built by manual docking in UCSF Chimera-1.19^150^. Refinement of the initial model was performed using real-space refine under Phenix2.0^151^. The OLD tetramer were rigid-body refined against the map, followed by real-space refinement with geometry, rotamer, Ramachan-dran-plot, Cβ, non-crystallographic-symmetry, secondary-structure, and reference-structure (initial model as reference) restraints, followed by global minimization and local-rotamer fitting. Secondary-structure annotation was inspected and edited using UCSF Chimera. OLD tetramer were subjected to iterative cycles of model building and refinement in Coot-0.9.8.1^152^. The final atomic model at was deposited in the Electron Microscopy Data Bank (EMDB) and the Protein Data Bank (PDB) with accession codes EMDB-69186 and PDB 23RJ (Table S1).

### tRNA cleavage assay

Assays of tRNA^Thr^ cleavage by OLD were carried out as described in a previous report^48^. Chemically synthesized oligonucleotide tRNA^Thr^ (1 μM, final concentration, Sangon Biotech) and OLD or its derivatives (tetramer, 0.8 μM, final concentration) were mixed with or without Oad2 (monomer, 12.8 μM, final concentration) in the reaction buffer (20 mM Tris-HCl, pH 7.6 and 50 mM KCl). Each sample was incubated at 37℃ for 90 min and then analyzed on a 6% polyacrylamide gel (acrylamide/bisacrylamide ratio of 29:1) in TBE (90 mM Tris, 44 mM boric acid, 2 mM EDTA) followed by SYBR-Gold staining. tRNA bands were visualized using a gel documentation system (Tanon 2500R).

### ChIP-seq with OLD in *ΔrecB* cells

*ΔrecB* BW25113 was grown in LB in the presence of kanamycin to OD=0.4, washed with water, electroporated with pBR_OLD_E402A_3XFlag plasmid, and grown on LB-agar plate with tetracycline. Single colonies were grown in LB in the presence of kanamycin and tetracycline.

For ChIP-seq experiment, overnight LB culture grown in the presence of kanamycin and tetracycline was diluted 1:100 with LB without antibiotics (total volume 600 mL) and grown to OD=0.6, followed by adding 16% formaldehyde to 0.25%. Cells were collected by centrifugation 5 minutes at 5000g and re-suspended in 25mL Lysis buffer 3’ [10mM Tris pH8.0; 100mM NaCl; 1mM EDTA; 0.5mM EGTA] supplemented with human lysozyme 1mg/ml and protease inhibitor cocktail (Roche). Sample was sonicated in an ice-bath for 10 min with constant pulse at a midi-tip (power output 70%) in a metal beaker. The resulting lysate was spun 30 minutes at 30 000g and 20uL of the supernatant were taken as an input control. The rest of the lysate was mixed with 2500 units of EndoCleava (Benzonase, CCNBio) and incubated 1 hour on ice. 0.1% Sodium Deoxycholate; 0.5% N-lauroylsarcosine and 1% Triton X-100 were added to the sample before 20ug anti-FLAG antibodies were added as well. Sample was rotated overnight at +4C in 50mL Falcone tubes.

Protein A/G magnetic beads (100μl - dry bed, Pierce) pre-equilibrated with 1ml of ice cold 1X PBS+0.5% BSA (blocking solution) were added to the sample and incubated with rotation for 1 hour at +4C. Beads were separated on magnetic stand and transferred into fresh 1.7mL Eppendorf tube in 1ml of RIPA buffer [50 mM HEPES pH7.5; 250 mM LiCl; 1mM EDTA; 1% NP40; 0.7% Sodium Deoxycholate]. Beads were rotated for 10 minutes at +4C, separated on a magnetic stand and the wash was repeated for total of five times. Beads were washed once more in 1ml of TE buffer as above before reversal of crosslinked-sonicated-chromatin. Washed sample and input control were mixed with 250uL of TE buffer and incubated overnight at 65°C shaking with 1% SDS and 2 mg/mL proteinase K (from 20 mg/mL stock solution in water, GoldBio).

Next day samples were briefly spun down, beads separated on a magnetic stand and DNA was purified by using Chip DNA Clean and concentrator kit (Zymo-research). DNA concentration was measured by Qubit.

### ChIP-seq with OLD during λ phage infection

BW25113 was grown in LB to OD=0.4, washed with water, electroporated with pBR_OLD_E402A_3XFlag plasmid, and grown on LB-agar plate with tetracycline. Single colonies were grown in LB in the presence of tetracycline. For ChIP-seq experiment, overnight LB culture grown in the presence of 0.2% maltose, 10 mM MgSO4, and tetracycline was diluted 1:100 with LB supplemented with 0.2% maltose and 10 mM MgSO4 (total volume 100 mL) and grown to OD=0.9. Next, 10 mL lambda phage in LB (3.5*10^10), room temperature, was added to obtain MOI ∼5. Culture was mixed by brief swirling and allowed to grow further (during the first 3 min rotation was off). 34-mL aliquots were then taken into 16% formaldehyde to the final concentration of 0.25 % after 10, 15, or 25 min.

Cells were collected by centrifugation 5 minutes at 5000g and re-suspended in 1mL Lysis buffer each supplemented with human lysozyme 1mg/ml and protease inhibitor cocktail (Roche). Samples were sonicated for 10 minutes using Covaris sonicator. The resulting lysates were spun 10 minutes at 24 000g and 5uL of the supernatant were taken as an input control. 0.1% Sodium Deoxycholate; 0.5% N-lauroylsarcosine and 1% Triton X-100 were added to the samples before 5ug anti-FLAG antibodies were added as well. Samples were rotated overnight at +4C in Eppendorf tubes.

Protein A/G magnetic beads (100μl - dry bed, Pierce) pre-equilibrated with 1ml of ice cold 1X PBS+0.5% BSA were added to the sample and incubated with rotation for 1 hour at +4C. Beads were separated on magnetic stand in 1ml of RIPA buffer. Beads were rotated for 10 minutes at +4C, separated on a magnetic stand and the wash was repeated for total of five times. Beads were washed once more in 1ml of TE buffer as above before reversal of crosslinked-sonicated-chromatin. Washed sample and input control were mixed with 250uL of TE buffer and incubated overnight at 65°C shaking with 1% SDS and 2 mg/mL proteinase K (from 20 mg/mL stock solution in water, GoldBio).

Next day samples were briefly spun down, beads separated on a magnetic stand and DNA was purified by using Chip DNA Clean and concentrator kit (Zymo-research). DNA concentration was measured by Qubit.

### ChIP-seq with OLD in the presence of pKB2 plasmid

Wt BW25113 was grown in LB to OD=0.4, washed with water, co-electroporated with pBR_OLD_E402A_3XFlag and pKB2 plasmids, and grown on LB-agar plate with tetracycline and carbenicilin. Single colonies were grown in LB in the presence of tetracycline and carbenicilin.

For ChIP-seq experiment, overnight LB culture grown in the presence of tetracycline and carbenicillin was diluted 1:100 with LB without antibiotics (total volume 600 mL) and grown to OD=0.6, followed by adding 16% formaldehyde to 0.25%. The ChIP-seq was performed exactly as for *ΔrecB* BW25113 carrying pBR_OLD_E402A_3XFlag plasmid.

## Data and code availibility

The atomic model coordinates for P2 OLD tetramer have been submitted to the Protein Data Bank (https://www.rcsb.org/) with PDB ID 23RJ. Corresponding EM maps have been submitted to the Electron Microscopy Data Bank (https://www.ebi.ac.uk/pdbe/emdb/) with ID EMD-69186. CRISPRi sequencing data have been deposited to the European Bioinformatics Institute database under accession code PRJEB115325. Phage escapers sequencing data have been deposited in NCBI database under BioProject ID PRJNA1462654.

## Acknowledgments

We thank Gerald R Smith (Fred Hutchinson Cancer Center) for sharing λ phage mutants and the LLS phage, and Grzegorz Wegrzyn (University of Gdansk) for pKB2 plasmid.

## Funding

This work was supported by Russian Scientific Foundation (RSF) grants 24-74-10089 and 25-44-02137 to A.I. Rapiflex mass-spectrometry was performed at Advanced Mass-Spectrometry Core Facility. A.D. was supported by Helicon students grant. This work was supported by the Blavatnik Family Foundation (to E.N.) and the Howard Hughes Medical Institute (to E.N.). V.M. and M.T. were supported by the RSF grant 25-74-30007. This study was funded by European Research Council [101044479] to D.B., Agence Nationale de la Recherche [ANR-10-LABX-62-IBEID] to D.B. This work was supported by National Key Research and Development Program of China (2024YFC2309300) to C.W., and the Natural Science Foundation of China (32521004) to C.W.

## Authors Contribution

A. Isaev conceived the study; A. Derzhaev, and S.Belukhina performed *in vivo* experiments; J. Zhang, H.Song, and C. Wang obtained cryo-EM structure, and performed *in vitro* experiments together with V. Molodtsov, and M. Tikhomirova; A. Gavrilov, I. Shamovsky, V. Epshtein, and E. Nudler performed ChIP-seq experiments; A. Shenfeld identified Oad2 protein; F. Depardieu and D. Bikard performed CRISPRi experiment; B. Saudemont and D. Bikard performed RNA-sequencing; O. Burenina performed Northern blots; A. Demkina and K.Severinov performed phage mutants sequencing; M. Skutel assembled phage genomes; A. Isaev, C. Wang, K. Severinov, and E. Nudler secured resources for the study. A. Derzhaev, C. Wang, and A. Isaev wrote the manuscript. All authors reviewed, edited, and approved the final version.

## Declaration of interests

The authors declare no competing interests.

## Resource availability

Further information and requests for resources and reagents should be directed to and will be fulfilled by the lead contact, Artem Isaev.

## Materials availability

All unique bacterial strains, phages, and plasmids generated in this study are available from the lead contact without restriction.

## Declaration of generative AI and AI-assisted technologies in the manuscript preparation process

During the preparation of this work, the author(s) used DeepSeek to assist with grammar checking. The author(s) reviewed and edited the output as needed and take full responsibility for the content of the published article.

---

**Figure S1.**
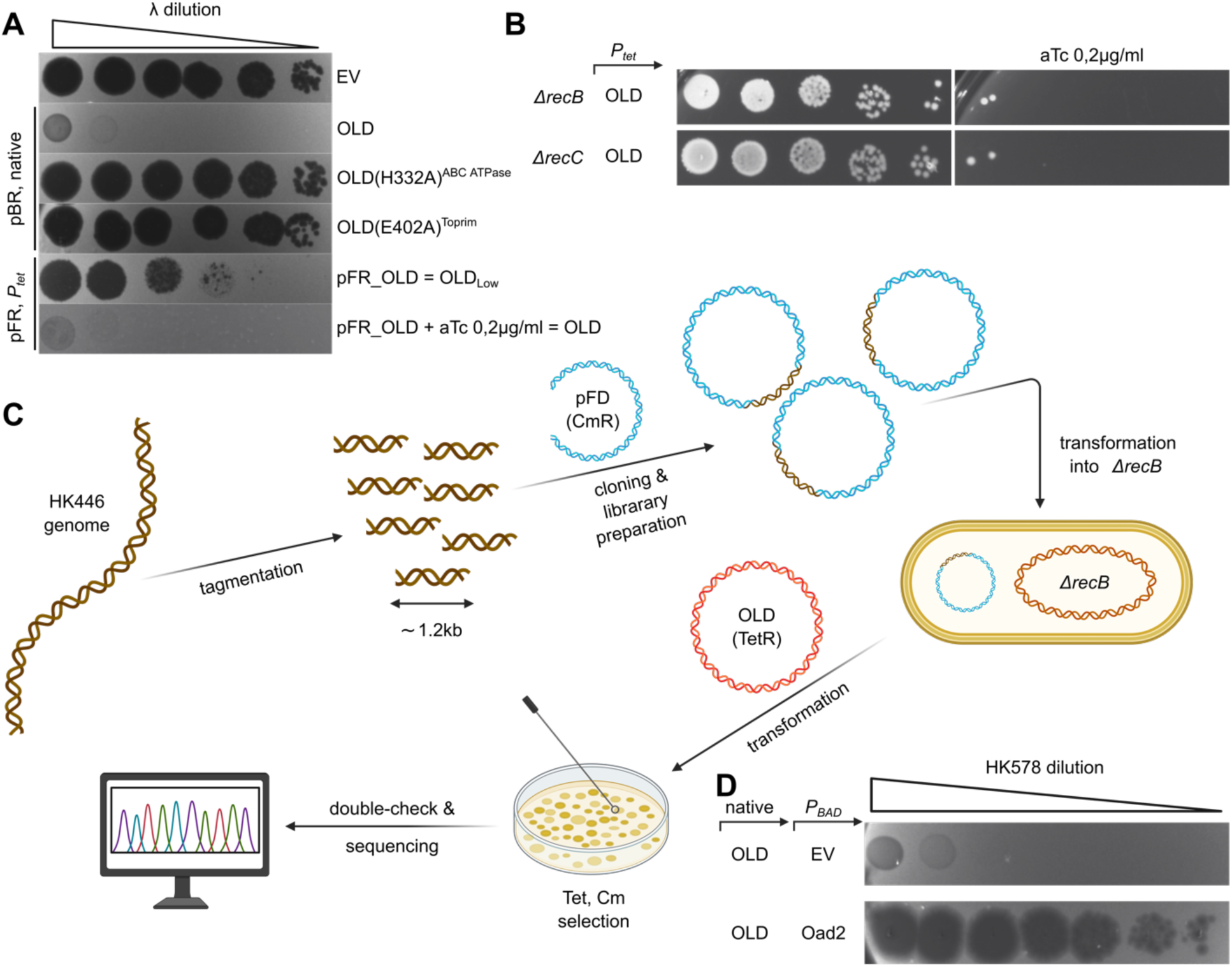
OLD cloning and mutagenesis, RecBCD-dependent toxicity, and discovery of the phage-encoded inhibitor Oad2. **(A)** EOP assay demonstrating that mutations in the Walker A motif (H332A) and the TOPRIM catalytic site (E402A) abolish OLD-mediated defense. Expression of OLD from its native promoter in pBR322 vector is compared to inducible expression from the pFR vector. Uninduced pFR_OLD (OLD_Low_) confers weak protection, while induction with anhydrotetracycline (aTc) restores defense to levels comparable to native expression. **(B)** Toxicity of inducible OLD in *ΔrecB* and *ΔrecC* strains, confirming that OLD activation is lethal in the absence of functional RecBCD. **(C)** Scheme of the plasmid library construction and screening strategy used to identify *Oad2* as an OLD inhibitor from the HK446 phage genome. **(D)** EOP assay showing that expression of Oad2 abolishes OLD-mediated defense against sensitive phage.

**Figure S2.**
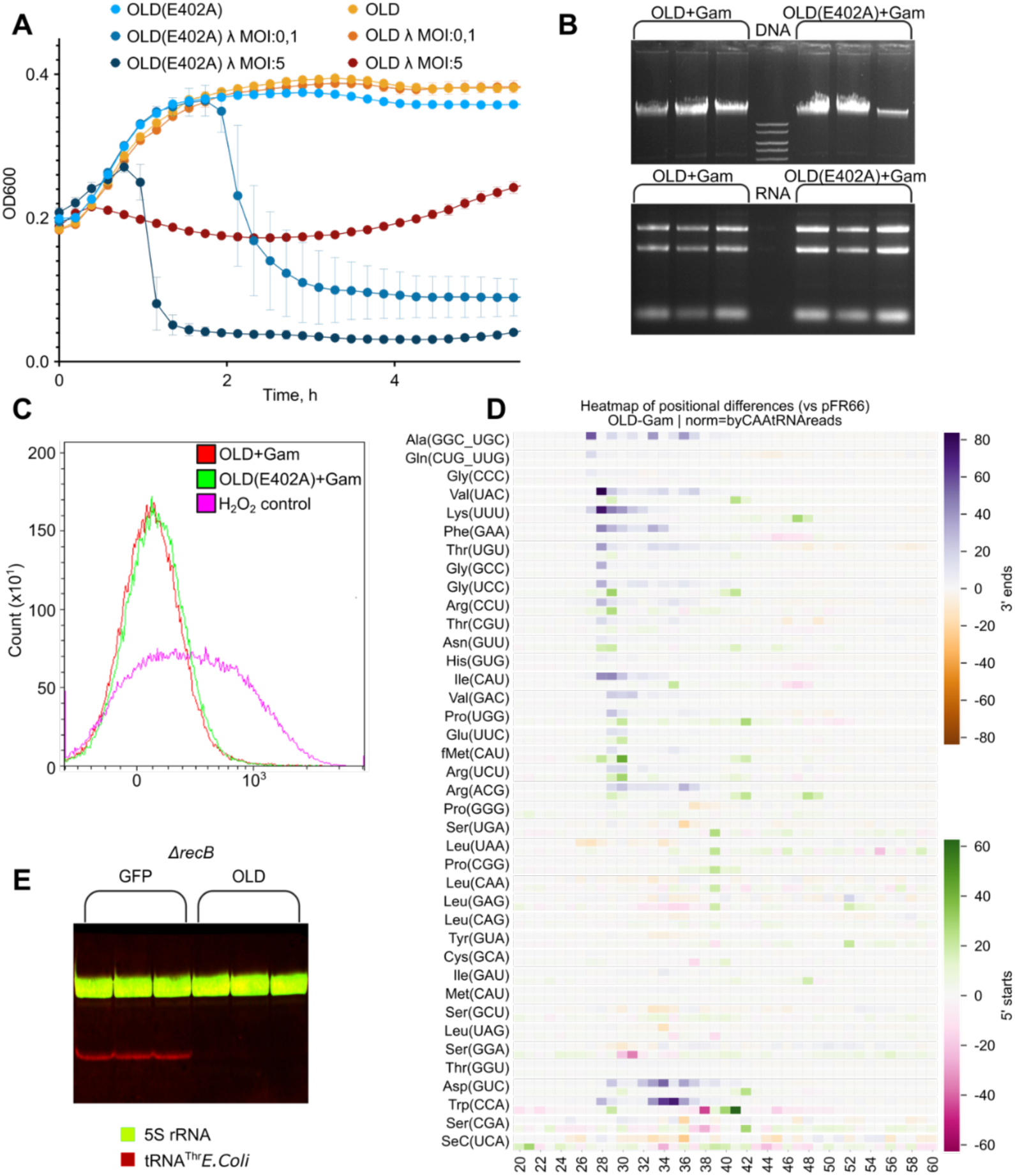
OLD induces abortive infection and cleaves host tRNAs without degrading DNA *in vivo*. **(A)** Growth curves of OLD-expressing cultures infected at low MOI (0.1) resemble uninfected growth, while control cultures lyse. At high MOI (5), OLD-expressing cultures arrest growth prior to lysis, confirming an abortive infection phenotype. **(B)** Agarose gel electrophoresis of total DNA and total RNA extracted from cells expressing OLD and Gam shows no detectable degradation compared to controls, indicating absence of general nucleic acid breakdown. **(C)** Terminal deoxynucleotidyl transferase dUTP nick end labeling (TUNEL) assay reveals no detectable DNA strand breaks in cells upon induction of Gam in OLD cells. **(D)** Heatmaps display differential 3ʹ-end (top) and 5ʹ-start (bottom) read accumulation between OLD and GFP samples (OLD − GFP), shown for positions 20–60 of each tRNA. Values were normalized using reads whose 3ʹ termini match the annotated mature tRNA end. Normalized differences were transformed using a signed log_10_ scale. The 3ʹ-end panel uses a purple-to-orange colormap (purple indicating enrichment in OLD), and the 5ʹ-start panel uses a green-to-pink colormap (green indicating enrichment in OLD). **(E)** Northern blot analysis of tRNA Thr(UGU) stability in *ΔrecB* cells expressing OLD or GFP control. Endogenous *E. coli* tRNA is degraded. Hybridization was performed step by step with *E. coli* anti- tRNA Cy3-labeled probe, then with *E. coli* anti- 5S rRNA probe (FITC).

**Figure S3.**
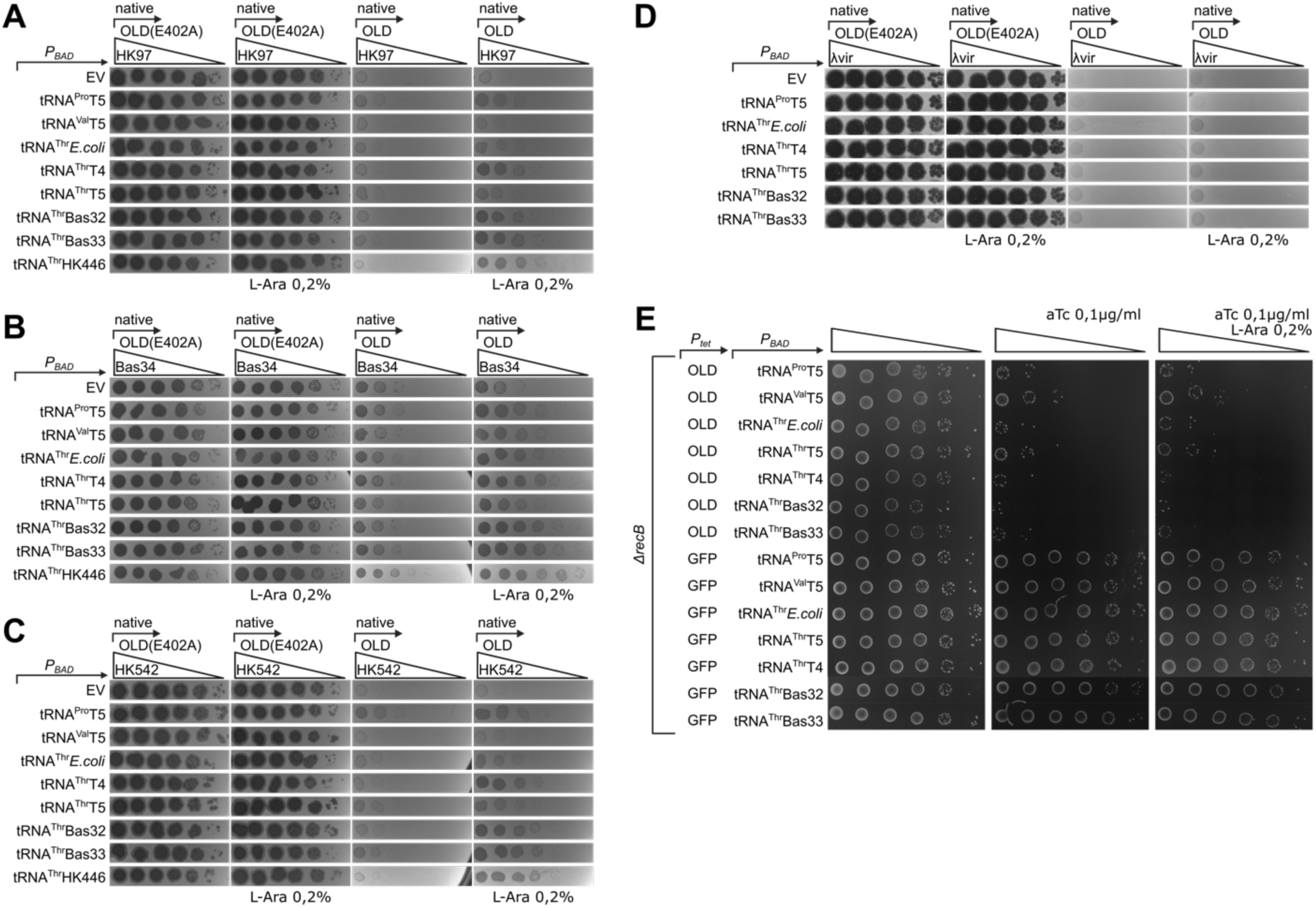
Viral tRNAs rescue HK97, Bas34, and HK542 phages from OLD defense, but are unable to rescue λ or inhibit OLD toxicity in *ΔrecB* cells. **(A, B, C, D)** Efficiency of plating (EOP) assays confirm that expression of phage-encoded tRNAs partially restores plaque formation of OLD-sensitive phages HK97 **(A)**, Bas34 **(B)**, HK542 **(C)**. Growth of phage λ **(D)** was not restored by tested tRNAs. **(E)** Toxicity assay of OLD expression in *ΔrecB* cells supplemented with phage-encoded tRNAs. Neither of tRNAs restored cell growth.

**Figure S4.**
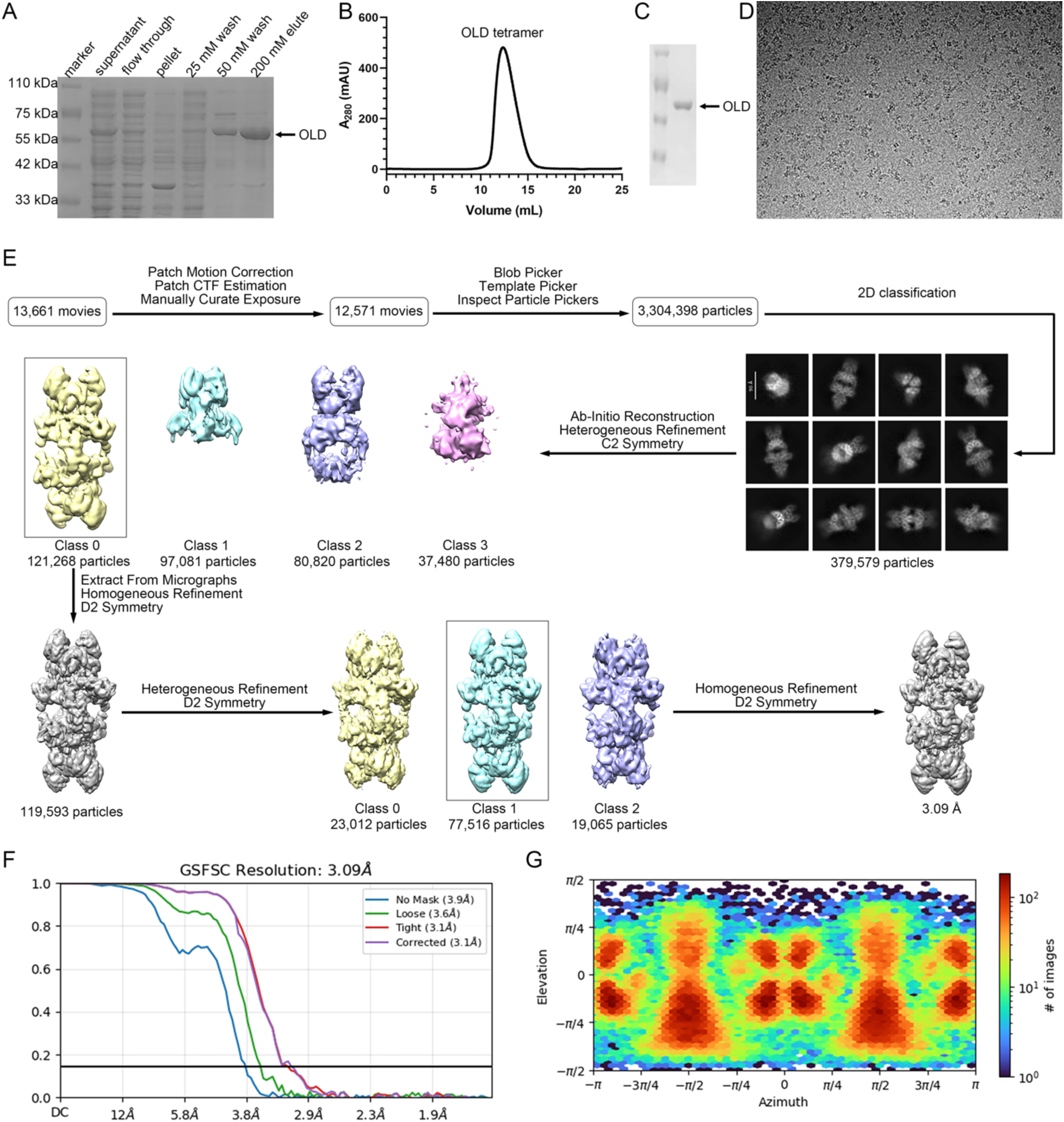
Structure analysis of P2 OLD. **(A)** Full-length OLD was expressed in *E. coli* BL21 (DE3) cells and was purified using Ni-NTA agarose. SDS-PAGE followed by Coomassie-blue staining showing the fractions collected at each step of purification with concentrations of imidazole labeled in elution part. **(B)** The purified OLD was subjected to size-exclusion chromatography (Superdex200 10/300). The molecular weight of OLD tetramer was about 264 kDa. **(C)** The protein purity visualized by SDS-PAGE. **(D)** Representative electron micrograph of the purified OLD. **(E)** Flow chart of cryo-EM data processing and 3D reconstruction of OLD. **(F)** FSC curve for the final reconstruction of the OLD tetramer. **(G)** The orientation distribution plot of the 3D reconstruction of OLD.

**Figure S5.**
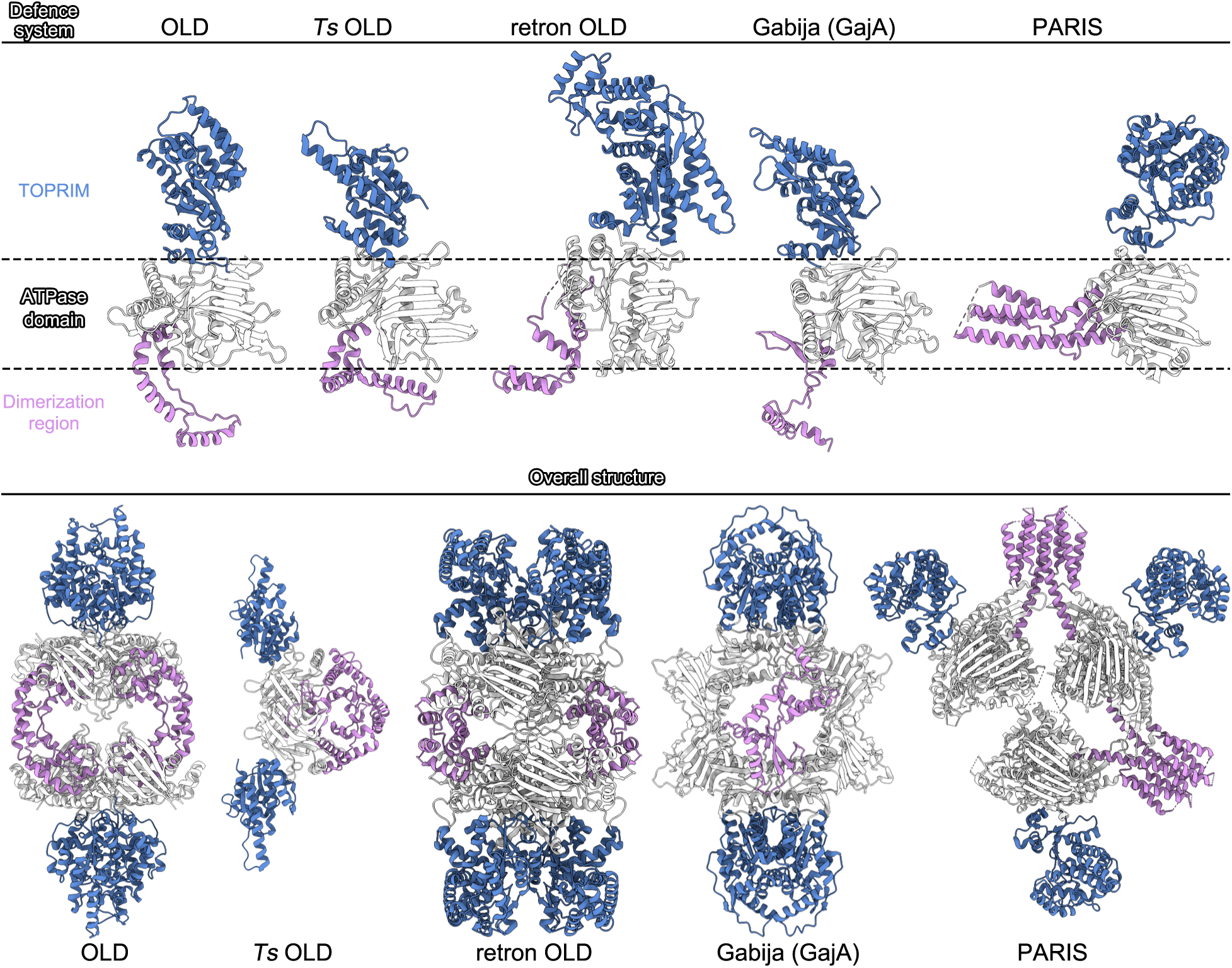
Structural comparison of OLD with other ABC+TOPRIM systems. Structural comparison of P2 OLD with *Thermus scotoductus* OLD (PDB ID: 6P74), retron OLD (9LP9), Gabija (GajA, PDB ID: 8X5I), and PARIS (8UX9). Upper panel: comparison between monomers; lower panel: comparison between oligomeric complexes.

**Figure S6.**
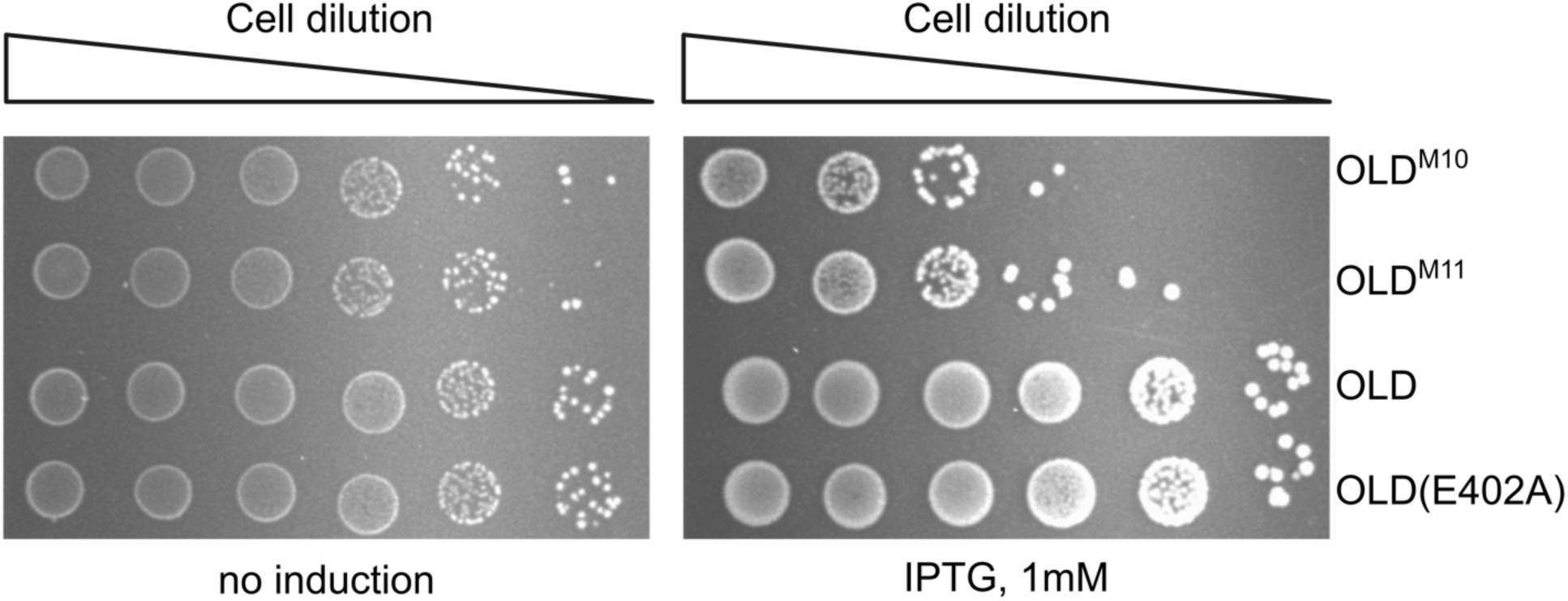
Dimeric OLD mutants are toxic to the host. Cell dilution assay demonstrating enhanced toxicity of pRSF_OLD^M10^ and pRSF_OLD^M11^ expression *in vivo*.

**Figure S7.**
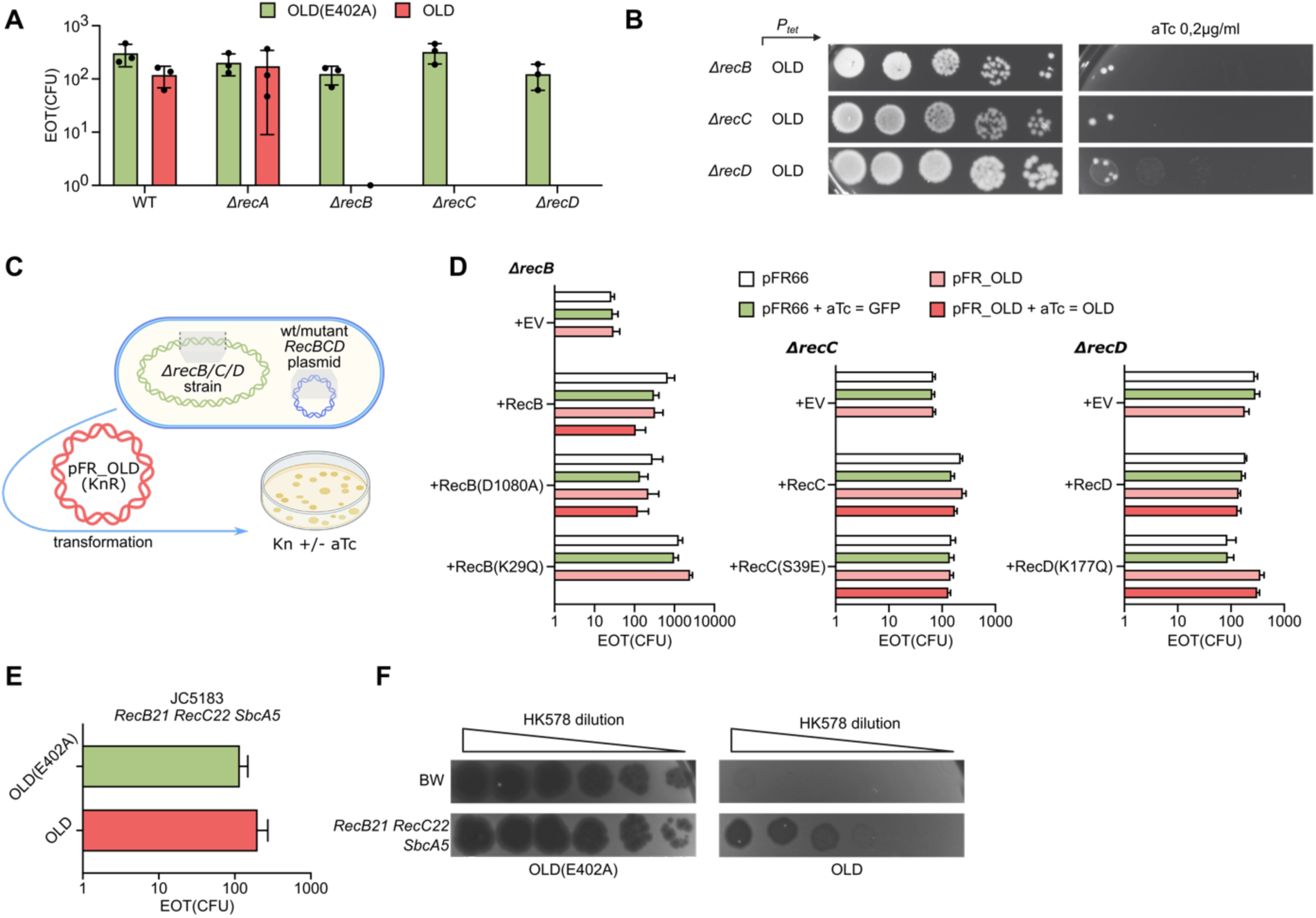
OLD toxicity is linked to RecBCD translocation deficiency and can be suppressed by alternative recombination pathways. **(A)** Efficiency of transformation (EOT) assay of the OLD-expressing plasmid in wild-type, Δ*recA*, Δ*recB*, Δ*recC*, and Δ*recD* strains. OLD is toxic specifically in Δ*recB*, Δ*recC*, and Δ*recD* backgrounds, but not in Δ*recA* or wild-type cells. **(B)** Toxicity of inducible OLD in *ΔrecB, ΔrecC* and *ΔrecD* strains, confirming that OLD activation is lethal in the absence of functional RecBCD. **(C)** Schematic of the EOT complementation assay. *ΔrecB, ΔrecC* or *ΔrecD* strains were first transformed with plasmids encoding either wild-type RecBCD or the indicated mutant variants, followed by transformation with pFR_OLD or the pFR66 control vector. Transformants were plated on selective medium supplemented with (+) or without (–) anhydrotetracycline (aTc) to induce OLD expression. **(D)** EOT assay demonstrating that OLD toxicity in Δ*recB*, Δ*recC*, or Δ*recD* strains is suppressed by complementation with wild-type RecBCD or with catalytically deficient mutants [RecB(D1080A), RecC(S39E), RecD(K177Q)], but not by the translocation-defective RecB(K29Q) mutant. Non-induced controls are shown for each condition. **(E)** EOT assay in the *E.coli* JC5183 (*recB21 recC22 sbcA5*) strain shows that activation of the RecET pathway suppresses OLD toxicity. **(F)** Efficiency of plating (EOP) assay demonstrates that OLD retains anti-phage activity in JC5183 cells, indicating phage-specific triggers remain effective even in *recB21 recC22 sbcA5* cells.

**Figure S8.**
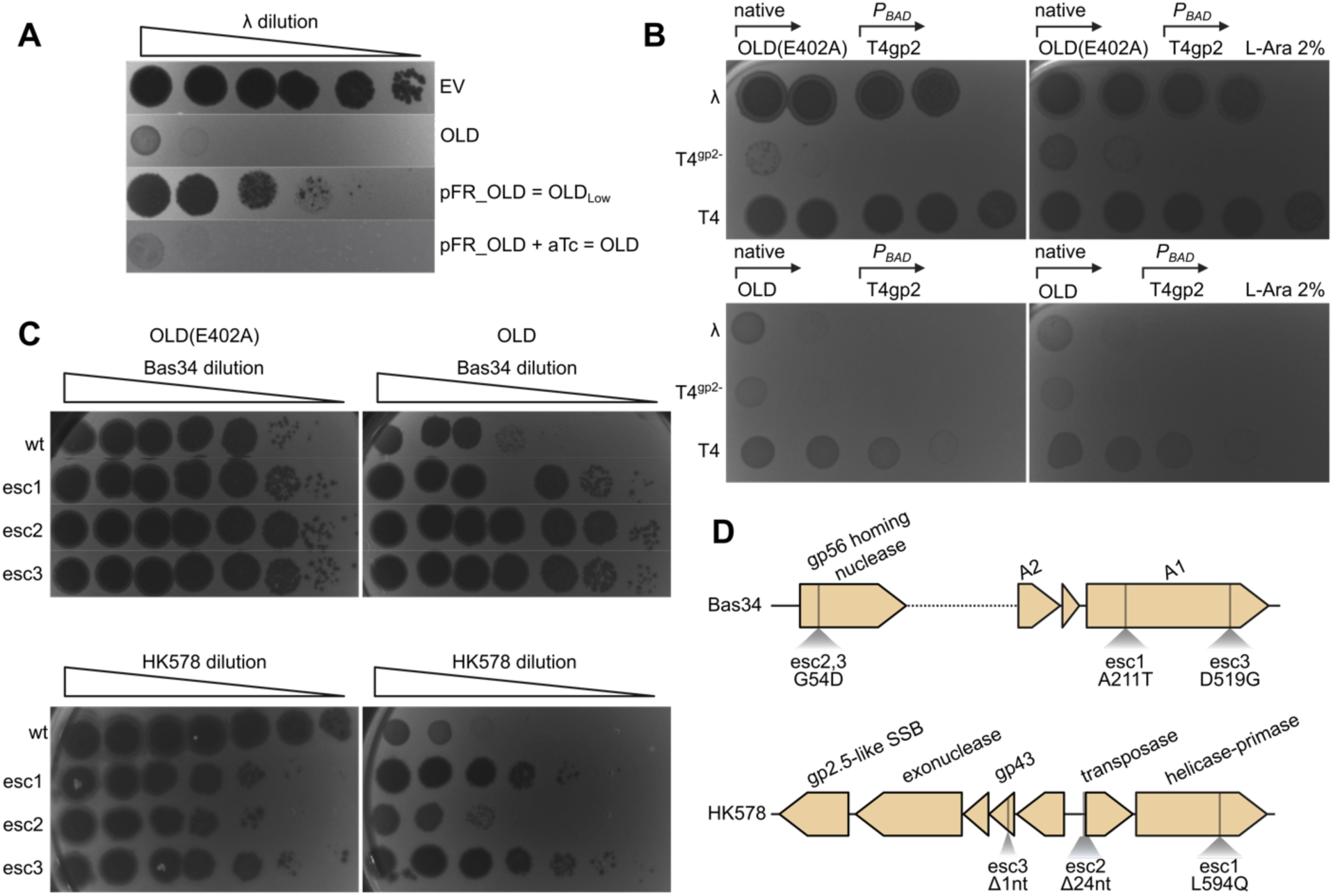
Identification of additional OLD phage escapers. **(A)** Efficiency of plating (EOP) assay comparing OLD expressed from its native promoter with inducible expression from the pFR vector, with and without anhydrotetracycline (aTc) induction. Native OLD confers full protection against λ, matching the protection level of induced pFR_OLD. Uninduced pFR_OLD shows only a slight EOP reduction and therefore is named OLD_Low_. **(B)** Expression of T4 gp2 does not rescue λ from OLD, indicating that OLD activation involves triggers beyond double-strand breaks (DSBs). **(C)** Schematic of Bas34 and HK578 escaper phage mutants, indicating the locations of mutations acquired in independent escapers. Bas34 acquired mutations in *a1* nuclease, or in *gp56* putative homing nuclease. HK578 escapers acquired mutations in replication initiation related genes (*gp46* helicase-primase, *gp45* promoter) or genes associated with DNA processing (*gp43*, adjacent to an exonuclease/SSB operon). **(D)** EOP assay of HK578 and Bas34 escaper mutants. HK578 esc2 exhibits only partial escape from OLD defense, while all other tested escapers are fully resistant.

**Figure S9.**
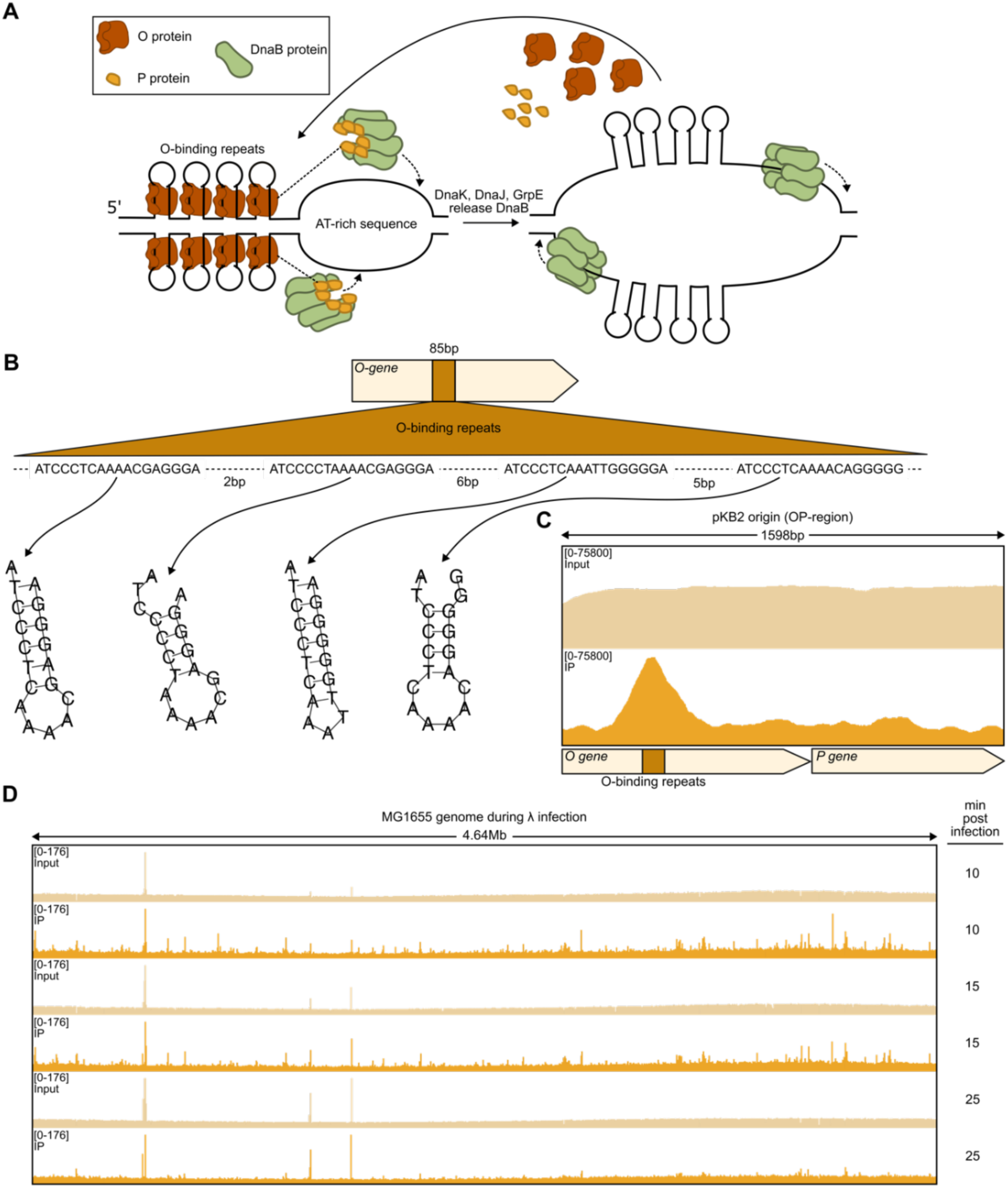
Model of the λ replication origin. OLD specifically binds the λ replication origin but not the chromosomal terminus during infection. **(A)** Model of λ origin DNA during theta replication, highlighting the formation of ssDNA hairpin structures at iterons within the melted region. **(B)** Schematic of the λ O gene and its binding region, composed of four iterons—palindromic repeats predicted to form ssDNA hairpins upon melting. **(C)** Chromatin immunoprecipitation sequencing (ChIP-seq) in cells harboring the pKB2 plasmid (*oriλ* + O/P genes) confirms specific OLD binding to the phage replication origin region *in vivo*. **(D)** Chromatin immunoprecipitation sequencing (ChIP-seq) during λ infection reveals no specific OLD binding to the terminus region, in contrast to what is observed in *recB*-deficient cells.

**Figure S10.**
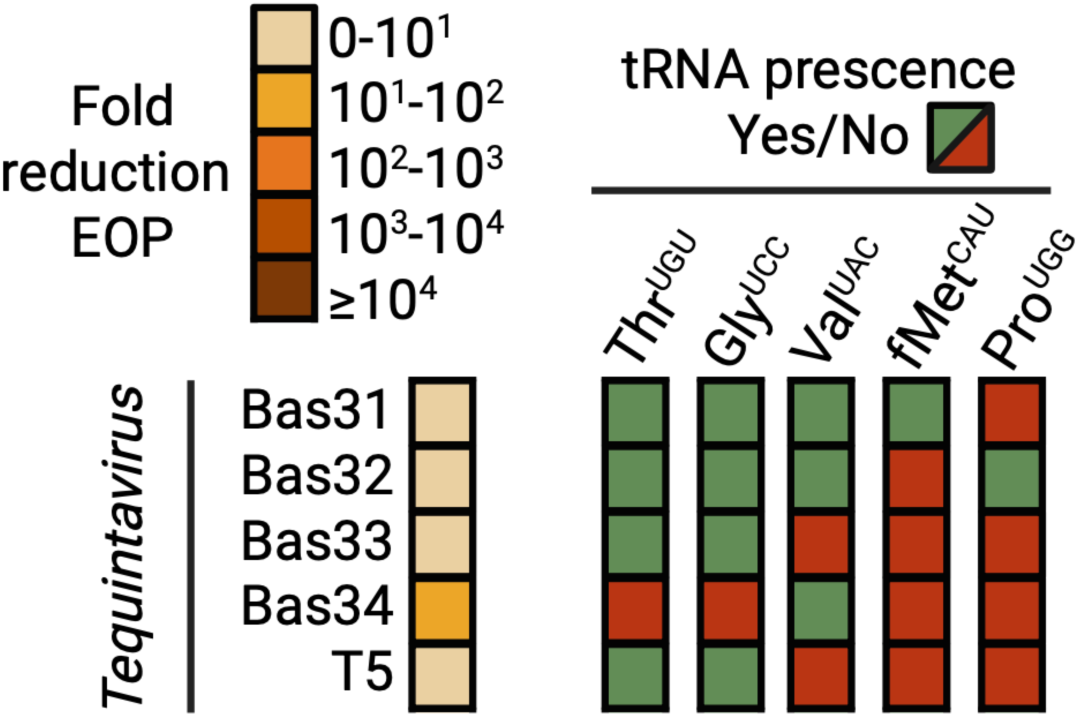
Presence of OLD-targeted tRNAs in *Tequintavirus* and relative phage defense. Correlation between the efficiency of OLD defense, evaluated via an EOP assay, and the presence of OLD-specific tRNA targets in *Tequintavirus* phages.

## Notes

### Competing Interest Statement

The authors have declared no competing interest.

