## Supplementary Tables. for "OLD sentinel: an abortive tRNase surveys phage replication and DNA defects in RecBCD-compromised cells"

### Table S1: Cryo-EM structure: P2-Old

| | **P2-OLD** (EMDB-69186) (PDB 23rj) |
|---|---|
| **Data collection and processing** | |
| Magnification | 105,000 |
| Voltage (kV) | 300 |
| Electron exposure (e–/Å$^2$) | 50 |
| Defocus range (μm) | -0.8 to -2.0 |
| Pixel size (Å) | 0.824 |
| Symmetry imposed | D2 |
| Initial particle images (no.) | 3,304,398 |
| Final particle images (no.) | 379,579 |
| Map resolution (Å) | 3.1 |
| FSC threshold | 0.143 |
| Map resolution range (Å) | 3.1-7.0 |
| **Refinement** | |
| Initial model used | Alphafold 2 |
| Model resolution (Å) | 3.1 |
| FSC threshold | 0.5 |
| Model composition | |
| Non-hydrogen atoms | 18392 |
| Protein residues | 2344 |
| Ligands | N/A |
| B factors (Å) | |
| Protein | 111.13 |
| Ligands | N/A |
| R.m.s. deviations | |
| Bond lengths (Å) | 0.003 |
| Bond angles ( ₒ) | 0.803 |
| Validation | |
| MolProbity score | 2.30 |
| Clashscore | 18.32 |
| Poor rotamers (%) | 0 |
| Ramachandran plot | |
| Favored (%) | 90.50 |
| Allowed (%) | 9.50 |
| Disallowed (%) | 0 |

### Table S2: Phages used in the study

| Name | Reference | Genome |
|---|---|---|
| Bas26 | [1] | NC_105113.1 |
| Bas27 | [1] | NC_105114.1 |
| Bas28 | [1] | NC_105115.1 |
| Bas29 | [1] | NC_105116.1 |
| Bas30 | [1] | NC_105117.1 |
| Bas31 | [1] | NC_105118.1 |
| Bas32 | [1] | NC_130053.1 |
| Bas33 | [1] | NC_105119.1 |
| Bas34 | [1] | MZ501103.1 |
| BF23 | [2] | OR083247.1 |
| DT57C | [3] | NC_027356.1 |
| Gostya9 | [4] | NC_047979.1 |

| | | |
|---|---|---|
| T5 | [5] | NC_005859.1 |
| MSK1 | Laboratory phage collection | N/A |
| MSK2 | Laboratory phage collection | N/A |
| MSK6 | Laboratory phage collection | N/A |
| MSK7 | Laboratory phage collection | N/A |
| HK97 | [6] | NC_002167.1 |
| HK446 | [6] | NC_019714.1 |
| HK544 | [6] | NC_019767.1 |
| HK75 | [6] | NC_016160.1 |
| HK106 | [6] | NC_019768.1 |
| HK542 | [6] | NC_019769.1 |
| HK140 | [6] | NC_019710.1 |
| HK243 | [6] | N/A |
| 9g | [7] | NC_024146.1 |
| mEp043 | [8] | NC_019706.1 |
| mEp213 | [8] | NC_019720.1 |
| HK578 | [6] | NC_019724.1 |
| MSK12 | Laboratory phage collection | N/A |
| MSK14 | Laboratory phage collection | N/A |
| lambda | [9] | NC_001416.1 |
| lambda872 | [10] | N/A |
| lambda1188 | [10] | N/A |
| lambda1191 | [10] | N/A |
| HK225 | [6] | NC_019717.1 |
| T6 | [5] | NC_054907.1 |
| T4 | [5] | NC_000866.4 |
| RB49 | [11] | NC_005066.1 |
| Brandy49 | [12] | MZ504876 |
| Cognac49 | [12] | MZ504877 |
| Whisky49 | [12] | MZ504878 |
| MSK8 | Laboratory phage collection | N/A |
| MSK10 | Laboratory phage collection | N/A |
| P2 | [13] | NC_001895.1 |
| P1 | [13] | NC_005856.1 |
| T7 | [5] | NC_001604.1 |
| lambda (ts), CmR | Laboratory phage collection | N/A |

### Table S3: Bacterial strains used in the study

| | | |
|---|---|---|
| *E. coli* Dh5a | Thermo Fisher (cat. 18265017) | NZ_CP076470.1 |
| *E. coli* BL21-AI | Thermo Fisher (cat. C607003) | NZ_CP047231.1 |
| *E. coli* K12 BW25113 | Laboratory strain collection | NZ_CP009273.1 |
| *E. coli* K-12 BW25113 r*ecB::kanR* | KEIO collection | |

| | | |
|---|---|---|
| *E. coli* K-12 BW25113 *recC::kanR* | KEIO collection | |
| *E. coli* K-12 BW25113 *recD::kanR* | KEIO collection | |
| *E. coli* K-12 BW25113 *sbcD::kanR* | KEIO collection | |
| *E. coli* K-12 BW25113 *ΔrecB* | KnR-depleted variant obtained from KEIO collection via pCPP20-mediated recombination | |
| *E. coli* K-12 BW25113 *ΔrecC* | KnR-depleted variant obtained from KEIO collection via pCPP20-mediated recombination | |
| *E. coli* K-12 BW25113 *ΔrecD* | KnR-depleted variant obtained from KEIO collection via pCPP20-mediated recombination | |
| *E. coli* JC5183 *F-, gal, recB21, recC22, sbcA5, endA* | [14] | N/A |
| *E. coli* MFDpir | [15] | |
| BL21(DE3) | Laboratory strain collection | |

### Table S4: Plasmids used in the study

| Name | Comment | Source |
|---|---|---|
| pFR66 | sfGFP, Ptet promoter, KanR | Bikard |
| pBR_OLD | OLD, native prromoter, TetR | This work |
| pBR_OLD(E402A) | OLD(E402A) TOPRIM mutant, native prromoter, TetR | This work |
| pBR_OLD(H332A) | OLD(H332A) ATPase mutant, native prromoter, TetR | This work |
| pFR_OLD | OLD, Ptet promoter, KanR | This work |
| pBAD_Gam | Gam λ, PBAD promoter, AmpR | This work |
| pBAD_gp5.9 | Gp5.9 λ, PBAD promoter, AmpR | This work |
| pBAD_reclocus_HK75 | rec locus HK75(gp33, Erf, gp31, Arf) AraBAD promoter, AmpR | This work |
| pBAD_gp33_HK75 | Gp33 HK75, AraBAD promoter, AmpR | This work |
| pBAD_Erf_HK75 | Gp33 HK75, AraBAD promoter, AmpR | This work |
| pBAD_gp31_HK75 | Gp33 HK75, AraBAD promoter, AmpR | This work |
| pBAD_Abc2_HK75 | Gp33 HK75, AraBAD promoter, AmpR | This work |
| pBAD_gp33_HK446 | rec locus HK446(gp33, Erf, gp31, Arf), PBAD promoter, AmpR | This work |
| pBAD_Oad2 | Oad2 (gp27 HK446), AraBAD promoter, AmpR | This work |
| pFD | Vector used for library preparation | [16] |
| pBAD_T5_tRNAPro | T5 tRNA Pro(UGG), AraBAD promoter, AmpR | [16] |
| pBAD_T5_tRNAVal | T5 tRNA Val(UAC), AraBAD promoter, AmpR | [16] |
| pBAD_E.coli_tRNAThr | E.coli tRNA Thr(UGU), AraBAD promoter, AmpR | This work |
| pBAD_T5_tRNAThr | T5 tRNA Thr(UGU), AraBAD promoter, AmpR | This work |

| | | |
|---|---|---|
| pBAD_T4_tRNAThr | T4 tRNA Thr(UGU), AraBAD promoter, AmpR | This work |
| pBAD_Bas32_tRNAThr | Bas32 tRNA Thr(UGU), AraBAD promoter, AmpR | This work |
| pBAD_Bas33_tRNAThr | Bas33 tRNA Thr(UGU), AraBAD promoter, AmpR | This work |
| pSA607 | RecBCD (RecC2773 6xHis), native promoter, AmpR | [17] |
| pSA335 | pSA607 derivative with RecB(D1080A), native promoter, AmpR | [17] |
| pSA618 | pSA607 derivative with RecB(K29Q), native promoter, AmpR | [17] |
| pJS24 | pSA607 derivative with RecC(S39E), native promoter, AmpR | [17] |
| pSA620 | pSA607 derivative with RecD(K177Q), native promoter, AmpR | [17] |
| pBAD_AddAB | AddAB B.subtillis, AraBAD promoter, AmpR | This work |
| pKB2_KanR | Plasmid with a λ replication origin, KanR | [18] |
| pBAD_T4gp2 | gp2 T4, PBAD promoter, AmpR | This work |
| pRSF_OLD | OLD(C-terminal 6xHis), T7lac promoter, KanR | This work |
| pRSF_OLD(E402A) | OLD(E402A)(C-terminal 6xHis), T7lac promoter, KanR | This work |
| pRSF_OLD_M10 | OLD_M10(C-terminal 6xHis), T7lac promoter, KanR | This work |
| pRSF_OLD_M11 | OLD_M11(C-terminal 6xHis), T7lac promoter, KanR | This work |
| pET22b_Oad2 | Oad2, T7lac promoter, AmpR | This work |
| pBAD_HK446_tRNAThr | HK446 tRNA Thr(UGU), AraBAD promoter, AmpR | This work |
| pKB2_AmpR | Plasmid with a λ replication origin, AmpR | [18] |
| pBR_OLD_E402A_3XFlag | OLD(E402A) TOPRIM mutant, C-terminal 3XFlag, native promoter, TetR | This work |
| pFR56 | dCas9, pPhlF promoter, CmR | [19] |
| pFR58 | pPhIF repressor, concstitutive promote, KnR | [19] |
| pFD265 | Gam, pPhIF promoter, CmR | This work |

### Table S5: Primers and other oligonucleotides used in the study

| Oligonucleotide name | Sequence (5'-3') | Template | Comment |
|---|---|---|---|
| pFR66_check_F | GAT AAA ACG AAA GGC CCA G | pFR_OLD | Sequencing primer |
| pFR66_check_R | AGC TCT CTA TCA TTG ATA GAG TG | pFR_OLD | Sequencing primer |
| OLD_pFR_F | GCC TTT CGT TTT ATT TGA TGC CTG GTT AAA TCC ATT TTA TGA AAT CTT CC | P2 phage | primer for OLD cloning into pFR |

| | | | |
|---|---|---|---|
| | | | vector via Gibson assembly |
| OLD_pFR_R | CAA ATG ACC TAG TTA GGA GGC AAA AAC TAA TGA GCC ATC AGT ATT TCC | P2 phage | primer for OLD cloning into pFR vector via Gibson assembly |
| pFR66_Ptet_F | TTT TGC CTC CTA ACT AGG TCA TTT G | pFR66 | primer for backbone amplification |
| pFR66_Ptet_R | CCA GGC ATC AAA TAA AAC GAA AGG | pFR66 | primer for backbone amplification |
| pBR_Old_F | CAT TAT TTT TCC TTA AAT GTG C | P2 phage | primer for OLD cloning into pBR vector via Gibson assembly |
| pBR_Old_R | ATA CAT TCA AAT ATG TAT CCG CTC AAC TAA TGA GCC ATC AGT ATT TCC | P2 phage | primer for OLD cloning into pBR vector via Gibson assembly |
| pBR322_Tetr_check_F | GAA AAA AAG GAT CTC AAG AAG ATC CT | pBR_OLD | Sequencing primer |
| pBR322_Tetr_check_R | CGG AAC CCC TAT TTG TTT ATT TTT C | pBR_OLD | Sequencing primer |
| pBR322_Tetr_F | TGA GCG GAT ACA TAT TTG AAT GTA T | pBR322 | primer for backbone amplification |
| pBR322_Tetr_R | AAG ATC CTT TTT GAT AAT CTC ATG AC | pBR322 | primer for backbone amplification |
| OLD_E402A_F | TGT TTT CCC TTC AAC AAG C | pBR_OLD | primer for OLD E402A mutagensis via KLD |
| OLD_E402A_R | GCG ACA AAC GTT CTA TAT GCA C | pBR_OLD | primer for OLD E402A mutagensis via KLD |
| OLD_H332A_F | AGT TGA TAT TAT AAC CTG ATA CC | pBR_OLD | primer for OLD H332A mutagensis via KLD |
| OLD_H332A_R | GCC TCA GCC AGT ATG CTT TC | pBR_OLD | primer for OLD H332A mutagensis via KLD |
| pBAD_Gam_F | TCG AGC TCT AAG GAG GTT ATA AAA AAT GGA TAT TAA TAC TGA AAC TGA G | λ phage | primer for Gam cloning into pBAD vector via Gibson assembly |

| | | | |
|---|---|---|---|
| pBAD_Gam_R | CAA GCT TGC ATG CCT GCA GGT CGA CTT ATA CCT CTG AAT CAA TAT CAA C | λ phage | primer for Gam cloning into pBAD vector via Gibson assembly |
| pBAD_gp5.9_F | TTT TTG GGC TAA CAG GAG GAA GAA TAT GTC TCG TGA CCT TGT GAC | T7 phage | primer for gp5.9 cloning into pBAD vector via Gibson assembly |
| pBAD_gp5.9_R | AAG CTT GCG GCC GCG AGC TCC ATG ATC AAG AAG TGC CAT TAA GTT TC | T7 phage | primer for gp5.9 cloning into pBAD vector via Gibson assembly |
| pBAD_gp33_HK75_F | TCG AGC TCT AAG GAG GTT ATA AAA AAT GAG TTT TAC AGA TAA CTG GTC | HK75 phage and HK446 phage | primer for gp33 or rec locus cloning into pBAD vector via Gibson assembly |
| pBAD_gp33_HK75_R | CAA GCT TGC ATG CCT GCA GGT CGA CTC ACG ATC TCC CTT CTG | HK75 phage | primer for gp33 cloning into pBAD vector via Gibson assembly |
| pBAD_Erf_HK75_F | TCG AGC TCT AAG GAG GTT ATA AAA AAT GGA TTT GAA TAA ATT CGA TGA G | HK75 phage | primer for Erf cloning into pBAD vector via Gibson assembly |
| pBAD_Erf_HK75_R | CAA GCT TGC ATG CCT GCA GGT CGA CTT ATG CCG CCT GTT TTA G | HK75 phage | primer for Erf cloning into pBAD vector via Gibson assembly |
| pBAD_gp31_HK75_F | TCG AGC TCT AAG GAG GTT ATA AAA AAT GGC AAG CAG AGG CGT AAA TAA | HK75 phage | primer for gp31 cloning into pBAD vector via Gibson assembly |
| pBAD_gp31_HK75_R | CAA GCT TGC ATG CCT GCA GGT CGA CTT AAA TTG CGT GAA TAG CGT GAC | HK75 phage | primer for gp31 cloning into pBAD vector via Gibson assembly |
| pBAD_Abc2_HK75_F | TCG AGC TCT AAG GAG GTT ATA AAA AAT GCC AGC GCC TCT GTA TGG | HK75 phage | primer for Abc2 cloming into pBAD vector via Gibson assembly |

| | | | |
|---|---|---|---|
| pBAD_Abc2_HK75_R | CAA GCT TGC ATG CCT GCA GGT CGA CTC AGT CAT TAC TGA TAG CGC CAT AG | HK75 phage and HK446 phage | primer for Abc2 or rec locus cloning into pBAD vector via Gibson assembly |
| 256_gp27_HK446_F | TCG AGC TCT AAG GAG GTT ATA AAA AAT GAA ACA AAT GAC ACT AAT TGA GAT G | HK446 phage | primer for Oad2 (gp27) cloning into pBAD vector via Gibson assembly |
| 256_gp27_HK446_R | CAA GCT TGC ATG CCT GCA GGT CGA CTC ACT GGT TGC CTC CTT TG | HK446 phage | primer for Oad2 (gp27) cloning into pBAD vector via Gibson assembly |
| pFD_Lin_F | CTG TCT CTT ATA CAC ATC TCC TGT CGG GTA GCA CCA GAA GTC | pFD | primer for backbone amplification |
| pFD_Lin_R | CTG TCT CTT ATA CAC ATC TCC TTT TTG CCA CCT GCA CTC GTC C | pFD | primer for backbone amplification |
| pFD_check_F | TCG TTT TGG TCA CGA CGT AC | pFD | Sequencing primer |
| pFD_check_R | AAT CCG GCA AGG AAA CAC T | pFD | Sequencing primer |
| pBAD_Ecoli_Thr_F | GGG CTA GCG AAT TCG AGC TCT CCA TAG AAT GCG CGC | E. coli | primer for Thr(UGU) cloning into pBAD vector via Gibson assembly |
| pBAD_Ecoli_Thr_R | TTG CAT GCC TGC AGG TCG ACG AAC CCC ACC GGA CTT G | E. coli | primer for Thr(UGU) cloning into pBAD vector via Gibson assembly |
| pBAD_T4_Thr_F | TTG CAT GCC TGC AGG TCG ACG CTG CTA ACC ATT GAG C | T4 phage | primer for Thr(UGU) cloning into pBAD vector via Gibson assembly |
| pBAD_T4_Thr_R | GGG CTA GCG AAT TCG AGC TCG GTT CAA ATC CTA TCG CCT C | T4 phage | primer for Thr(UGU) cloning into pBAD vector via Gibson assembly |
| pBAD_Bas32_Thr_F | GGG CTA GCG AAT TCG AGC TCG TTG AAA TAA CGG GAT TAT C | Bas32 phage | primer for Thr(UGU) cloning into pBAD vector via Gibson assembly |

| | | | |
|---|---|---|---|
| pBAD_Bas32_Thr_R | TTG CAT GCC TGC AGG TCG ACT GAA TTT TTA AAT TTG GTG C | Bas32 phage | primer for Thr(UGU) cloning into pBAD vector via Gibson assembly |
| pBAD_Bas33_Thr_F | GGG CTA GCG AAT TCG AGC TCC GTT GAA ATA ACG GGT TC | Bas33 phage | primer for Thr(UGU) cloning into pBAD vector via Gibson assembly |
| pBAD_Bas33_Thr_R | TTG CAT GCC TGC AGG TCG ACT AAC AGT TTA AAC ATA ATA GTC C | Bas33 phage | primer for Thr(UGU) cloning into pBAD vector via Gibson assembly |
| pBAD_HK446_Thr_F | TCG AGC TCT AAG GAG GTT ATA AAA AGG TTT CGG GAT TTT TTA TTT GGG TC | HK446 phage | primer for Thr(UGU) cloning into pBAD vector via Gibson assembly |
| pBAD_HK446_Thr_R | CAA GCT TGC ATG CCT GCA GGT CGA CTC ACT GAT GAG TTC AGG ATA GC | HK446 phage | primer for Thr(UGU) cloning into pBAD vector via Gibson assembly |
| 256b_F | TTT TTA TAA CCT CCT TAG AGC TCG | pBAD | primer for backbone amplification |
| 256b_R | GTC GAC CTG CAG GCA TGC | pBAD | primer for backbone amplification |
| pBAD_for | ATG CCA TAG CAT TTT TAT CC | pBAD | Sequencing primer |
| pBAD_rev | GAT TTA ATC TGT ATC AGG CTG | pBAD | Sequencing primer |
| pBAD_T5_Thr_F | GGGCTAGCGAATTCGAGCTCgttgaaataacgggatc | T5 phage | primer for Thr(UGU) cloning into pBAD vector via Gibson assembly |
| pBAD_T5_Thr_R | TTGCATGCCTGCAGGTCGACgcaaatgaatttttaag | T5 phage | primer for Thr(UGU) cloning into pBAD vector via Gibson assembly |
| pBAD_AddAB_F | TCG AGC TCT AAG GAG GTT ATA AAA AAT GGG AGC AGA GTT TTT AGT AGG | B. subtillis | primer for AddAB cloning into pBAD vector via Gibson assembly |

| | | | |
|---|---|---|---|
| pBAD_AddAB_R | CAA GCT TGC ATG CCT GCA GGT CGA CCT ATA ATG TCA GAA TGT GCC CTC | B. subtillis | primer for AddAB cloning into pBAD vector via Gibson assembly |
| pBAD_gp2_T4_F | TCG AGC TCT AAG GAG GTT ATA AAA AAT GGC TAT TTT TCA AAT AAT TAA TG | T4 phage | primer for gp2 cloning into pBAD vector via Gibson assembly |
| pBAD_gp2_T4_R | CAA GCT TGC ATG CCT GCA GGT CGA CTT AGA TAT TTG ACC AGA CTT TG | T4 phage | primer for gp2 cloning into pBAD vector via Gibson assembly |
| 5Phos_METagA1 | /5Phos/CTGTCTCTTATACACATCT | | Transposase oligo |
| METagA2 | AGATGTGTATAAGAGACAG | | Transposase oligo |
| E coli tRNA Thr Cy3 | /Cy3/- TAA TCA GTA GGT CAC CAG TTC HAT TCC GGT AGT CGG CAC CA | | For northern blot |
| Bas32 tRNA Thr Cy5 | /Cy5/-gCTCCCACCAggAATCgAACCCggTTCAgATgCTTACAAggC | | For northern blot |
| pRSF_OLD_F | TA CCATGGGC ACTGTACGTCTTGCTTCAGTTTCA | pBR_OLD | primer for OLD cloning into pRSF vector via NcoI restriction |
| pRSF_OLD_R | TA GCGGCCGC TTAGTGGTGATGATGGTGATGACCGCTACCAATCCATTTTATGAAATC | pBR_OLD | primer for OLD cloning into pRSF vector via NcoI restriction |
| pET22b_Oad2_F | ACTTTAAGAAGGAGATATACATATGAAACAAATGACACTAATTGAGAT | pBAD_Oad2 | primer for Oad2 climing into pET22b vector via Gibson assembly |
| pET22b_Oad2_R | ATCTCAGTGGTGGTGGTGGTGGTGCTGGTTGCCTCCTTTGCGAAGC | pBAD_Oad2 | primer for Oad2 climing into pET22b vector via Gibson assembly |
| pBR322_Old_3xFlag | TCA TGA GAT TAT CAA AAA GGA TCT TTT ACT TGT CAT CGT CAT CCT TGT AAT CGA TGT CAT GAT CTT TAT AAT CAC CGT CAT GGT CTT TGT AGT CAA TCC ATT TTA TGA AAT CTT CC | pBR_OLD(E402A) | Primer for FLAG-tag introduction via KLD |
| FR230 | TTCCCTACACGACGCTCTTCCGATCTTGCTNNNNGCACGCCCGTCGCTCAGTCCTAGGTATAATACTA | | Index 1 for custom sample preparation |
| FR231 | TTCCCTACACGACGCTCTTCCGATCTCTAANNNNGCACGCCCGTCGCTCAGTCCTAGGTATAATACTA | | Index 1 for custom sample preparation |

| | | | |
|---|---|---|---|
| FR232 | TTCCCTACACGACGCTCTTCCGATCTAGAGNNNNGC ACGCCCGTCGCTCAGTCCTAGGTATAATACTA | | Index 1 for custom sample preparation |
| FR233 | TTCCCTACACGACGCTCTTCCGATCTACTGNNNNGC ACGCCCGTCGCTCAGTCCTAGGTATAATACTA | | Index 1 for custom sample preparation |
| FR234 | TTCCCTACACGACGCTCTTCCGATCTCGTCNNNNGC ACGCCCGTCGCTCAGTCCTAGGTATAATACTA | | Index 1 for custom sample preparation |
| LC863 | GTGACTGGAGTTCAGACGTGTGCTCTTCCGATCTNN NNNNAAAGGACCCGTAAAGTGATAATGAT | | Reverse primer for sequencing PCR1 |
| LC1516 | CAAGCAGAAGACGGCATACGAGATGTCGTGATGTGA CTGGAGTTCAGACG | | Index 2 for custom sample preparation |
| LC415 | AATGATACGGCGACCACCGAGATCTACACTCTTTCCC TACACGACGCT | | Forward primer for sequencing PCR2 |
| LC609 | GCACGCCCGTCGCTCAGTCCTAGGTATAATACTA | | Custom read 1 primer for sequencing |
| LC499 | GATCGGAAGAGCACACGTCTGAACTCCAGTCAC | | Index primer 1 for sequencing |
| LC610 | TATTATACCTAGGACTGAGCGACGGGCGTGC | | Index primer 2 for sequencing |

1. Maffei, E., Shaidullina, A., Burkolter, M., Heyer, Y., Estermann, F., Druelle, V., Sauer, P., Willi, L., Michaelis, S., and Hilbi, H. (2021). Systematic exploration of Escherichia coli phage–host interactions with the BASEL phage collection. PLoS Biol. *19*, e3001424.
2. (Paris, S. de biologie (1918). Comptes rendus des séances de la Société de biologie (Masson et Cie).
3. Golomidova, A.K., Kulikov, E.E., Prokhorov, N.S., Guerrero-Ferreira, R.C., Ksenzenko, V.N., Tarasyan, K.K., and Letarov, A. V (2015). Complete genome sequences of T5-related Escherichia coli bacteriophages DT57C and DT571/2 isolated from horse feces. Arch. Virol. *160*, 3133–3137.
4. Golomidova, A.K., Kulikov, E.E., Babenko, V. V, Ivanov, P.A., Prokhorov, N.S., and Letarov, A. V (2019). Escherichia coli bacteriophage Gostya9, representing a new species within the genus T5virus. Arch. Virol. *164*, 879–884.
5. Demerec, M., and Fano, U. (1945). Bacteriophage-resistant mutants in Escherichia coli. Genetics *30*, 119.
6. Dhillon, E.K., Dhillon, T.S., Lam, Y.Y., and Tsang, A.H. (1980). Temperate coliphages: classification and correlation with habitats. Appl. Environ. Microbiol. *39*, 1046–1053.
7. Kulikov, E.E., Golomidova, A.K., Letarova, M.A., Kostryukova, E.S., Zelenin, A.S., Prokhorov, N.S., and Letarov, A. V (2014). Genomic sequencing and biological characteristics of a novel Escherichia coli bacteriophage 9g, a putative representative of a new Siphoviridae genus. Viruses *6*, 5077–5092.
8. Kameyama, L., Fernández, L., Calderón, J., Ortiz-Rojas, A., and Patterson, T.A. (1999). Characterization of wild lambdoid bacteriophages: detection of a wide distribution of phage immunity groups and identification of a Nus-dependent, nonlambdoid phage group. Virology *263*, 100–111.
9. Lederberg, E.M., and Lederberg, J. (1953). Genetic studies of lysogenicity in Escherichia coli. Genetics *38*, 51.

10. Amundsen, S.K., Sharp, J.W., and Smith, G.R. (2016). RecBCD enzyme "Chi recognition" mutants recognize Chi recombination hotspots in the right DNA context. Genetics *204*, 139–152.
11. Monod, C., Repoila, F., Kutateladze, M., Tétart, F., and Krisch, H.M. (1997). The genome of the pseudo T-even bacteriophages, a diverse group that resembles T4. J. Mol. Biol. *267*, 237–249.
12. Efimov, A.D., Golomidova, A.K., Kulikov, E.E., Belalov, I.S., Ivanov, P.A., and Letarov, A. V (2022). RB49-like bacteriophages recognize O antigens as one of the alternative primary receptors. Int. J. Mol. Sci. *23*, 11329.
13. Bertani, G. (1951). Studies on lysogenesis I: the mode of phage liberation by lysogenic Escherichia coli. J. Bacteriol. *62*, 293–300.
14. Barbour, S.D., Nagaishi, H., Templin, A., and Clark, A.J. (1970). Biochemical and genetic studies of recombination proficiency in Escherichia coli, II. Rec+ revertants caused by indirect suppression of Rec-mutations. Proceedings of the National Academy of Sciences *67*, 128–135.
15. Ferrieres, L., Hémery, G., Nham, T., Guérout, A.-M., Mazel, D., Beloin, C., and Ghigo, J.-M. (2010). Silent mischief: bacteriophage Mu insertions contaminate products of Escherichia coli random mutagenesis performed using suicidal transposon delivery plasmids mobilized by broad-host-range RP4 conjugative machinery. J. Bacteriol. *192*, 6418–6427.
16. Burman, N., Belukhina, S., Depardieu, F., Wilkinson, R.A., Skutel, M., Santiago-Frangos, A., Graham, A.B., Livenskyi, A., Chechenina, A., Morozova, N., et al. (2024). A virally encoded tRNA neutralizes the PARIS antiviral defence system. Nature *634*, 424–431. https://doi.org/10.1038/s41586-024-07874-3.
17. Amundsen, S.K., and Smith, G.R. (2019). The RecB helicase-nuclease tether mediates Chi hotspot control of RecBCD enzyme. Nucleic Acids Res. *47*, 197–209.
18. Baranska, S. (2002). Directionality of lambda plasmid DNA replication carried out by the heritable replication complex. Nucleic Acids Res. *30*, 1176–1181. https://doi.org/10.1093/nar/30.5.1176.
19. Rousset, F., Cabezas-Caballero, J., Piastra-Facon, F., Fernández-Rodríguez, J., Clermont, O., Denamur, E., Rocha, E.P.C., and Bikard, D. (2021). The impact of genetic diversity on gene essentiality within the Escherichia coli species. Nat. Microbiol. *6*, 301–312.
